# Species richness and trait diversity show parallel island-biogeographic patterns across Australian islands

**DOI:** 10.64898/2026.08.05.743155

**Authors:** Corey J. A. Bradshaw, Ashlea Naglis, Frédérik Saltré, Caitlin Mudge, Céline Bellard, Giovanni Strona, Vera Weisbecker, April E. Reside, Christopher R. Dickman, John Llewelyn

## Abstract

Island biogeography is the theoretical and empirically validated expectation that an island’s biodiversity is ultimately limited by its size and isolation, with larger and less isolated (from mainland communities) islands supporting greater diversity. Island biogeography is generally applied to simple metrics of diversity such as species richness (number of different species); however, fewer studies have used traits to measure biodiversity. An organism’s traits — e.g., body mass, age at sexual maturity, trophic level — can be used to measure biodiversity and understand how ecological communities function. We quantified species richness for birds, mammals, reptiles, and amphibians, and functional diversity for birds and mammals (for which sufficient trait data were available), and tested whether this diversity can be predicted using the theory of island biogeography. We identified 9,103 Australian islands, of which 1,661 had at least one (native and/or non-native) non-marine species present according to the Atlas of Living Australia. As expected, tetrapod species richness (*S*) increased with island area (*A*) (*z* = 0.299 ± 0.012) following a typical power-law relationship (i.e., *S* = *cA^z^*, where *z* = 0.2–0.4), but was not predicted by distance from mainland — consistent with the pattern observed on other recently (< 10,000 years) isolated continental islands. We found that trait richness increased at the same rate with island area as species richness for mammals, but for birds, trait richness increased more slowly than species richness. Trait turnover increased modestly with inter-island distance, whereas trait nestedness was unrelated to distance. Trait richness increased strongly with species richness in both birds and mammals, and island area and isolation explained no additional variation in functional richness after accounting for species richness. These results indicate that island geography influences functional diversity primarily through species accumulation, rather than through direct effects on occupied trait space. Overall, the trait space of smaller islands tended to be nested within that of larger, nearby islands; the main differences among similar-sized islands are due to turnover (change in species/trait combinations among assemblages), and the effects are more pronounced in mammals compared to birds. We also found evidence for an asymptotic relationship between trait and species turnover in both birds and mammals, suggesting close coupling between taxonomic and functional turnover, with some saturation of trait turnover at high species turnover. Large islands that are simultaneously more isolated might offer conservation advantages by reducing the influence of threatening processes on the mainland if distance limits access of people and invasive species.

## Introduction

Islands are disproportionately rich in biodiversity compared to similar-sized mainland areas, supporting an estimated 15–20% of terrestrial species despite comprising only 7% of Earth’s landmass (Gibson *et al*. 2017; Fernández-Palacios *et al*. 2021; Matthews & Triantis 2021). This unique richness, along with clearly defined boundaries, theoretically simpler ecosystems than mainland counterparts, and varying isolation from mainland communities, have made islands central to the development of ecological, evolutionary, and conservation theory (Matthews & Triantis 2021).

Based on the rates of immigration, evolution, and extinction of island species, island biogeography predicts a relationship between an island’s biodiversity and its size and isolation from the mainland and other islands (MacArthur & Wilson 1967). Accordingly, larger islands are predicted to have more biodiversity than smaller islands, and more isolated islands less biodiverse than those closer to the mainland (Matthews & Triantis 2021). Although there are many other determinants of total biodiversity on any given island (for example, an island’s age, elevation, location, primary production, and habitats), size and isolation are still considered some of the most influential determinants (Helmus & Behm 2020). Predictions of species richness can also be applied to ‘island’ patches of habitat within largely inhospitable matrices (e.g., woodland fragments in agricultural landscapes, colonising invertebrates on intertidal boulders, mountain-top communities) (Matthews & Triantis 2021; Matthews *et al*. 2026).

Species richness — the number of distinct species — has been the primary metric of biodiversity used in island-biogeographic analyses; however, the emerging field of trait-based ecology measures the ‘traits’ of organisms as another way of quantifying biodiversity patterns (Zakharova *et al*. 2019). Although definitions vary, a ‘trait’ is generally considered a measurable morphological, physiological, or behavioural characteristic that often relates to a species’ growth, reproduction, and survival (Violle *et al*. 2007). Body mass, trophic level, age at sexual maturity, and number of limbs are all examples of traits that inform how species function within their respective communities. The composition of traits across different species in an ecological community indicates how species interact and how the community functions as a whole (Llewelyn *et al*. 2025). Community trait composition, or ‘community trait space’, also sheds light on community-assembly rules, evolutionary dynamics, and how the composition of biodiversity might shift in the face of environmental change (Mouillot *et al*. 2021; Schrader 2026). To quantify how communities function and change over time, one can therefore measure different attributes of their trait space — including trait-space volume, location, and occupation density (Mouillot *et al*. 2021; Llewelyn *et al*. 2025). Because functional diversity metrics can correlate with species richness (Ricklefs & Miles 1994; Mammola *et al*. 2021), we hypothesise that trait diversity will also correlate positively with island area and negatively with island isolation from the mainland. However, these hypotheses have not yet been tested.

Australia’s ∼ 9,100 marine islands and islets provide an ideal system to test these hypotheses, because their areas vary widely, as do their distances from the mainland (Geoscience Australia 2023). Furthermore, many Australian islands have been surveyed for tetrapod taxa (including some with long-term survey records). Although there have been many studies in Australia that have tested the predictions of island biogeography on species composition in continental habitat patches — from viable weed seed diversity in pastures (Forcella 1984), plant species in heathlands (Keeley 2003) and eucalypt woodland patches (Morgan *et al*. 2011), invertebrates on stream stones (Douglas & Lake 1994), intertidal mussel colonies (Peake & Quinn 1993) and boulders (McGuinness 1984; Chapman & Underwood 2009), fish and mollusc assemblages in estuaries (Davis *et al*. 2017), birds in continental habitat patches (Maron *et al*. 2012; Storch *et al*. 2012), to lizards and non-volant mammals in forest reserves (Kitchener *et al*. 1980; Triantis *et al*. 2003) — only a few Australian studies have examined species-area relationships on small samples of true islands (Abbott & Black 1980; Buckley 1985; Woinarski *et al*. 1999a; Woinarski *et al*. 1999b; Woinarski *et al*. 2000; Lomolino 2001; Gao *et al*. 2019), or for a few taxa such as plants (Schrader *et al*. 2021) and amphibians/freshwater fishes (Ho *et al*. 2025) across a larger sample of Australian islands. Furthermore, only one study in Australia has examined the two main components of species and trait dissimilarity patterns — *turnover* (how species identities or trait combinations change among assemblages independently of differences in richness) and *nestedness* (extent to which species-poor or functionally depauperate assemblages contain subsets of the species, traits, or functional space present in richer assemblages) (Baselga 2010) — among islands in the context of island biogeography: a study of 156 plant species on 15 small and medium islands in Western Australia (Schrader *et al*. 2023). A broader examination among other taxa and across a wide range of island sizes and types has not yet been attempted, despite trait diversity being more functionally relevant than species richness (Schrader *et al*. 2021).

To test whether trait diversity follows similar patterns to species richness on true islands (including continental and oceanic islands) as a function of area and isolation, we obtained Australia-wide data describing species richness of birds, reptiles, amphibians, and mammals relative to island area and isolation. We then obtained available trait data for birds and mammals to see if similar patterns emerged. We predicted that smaller, more isolated islands will have lower species diversity (richness) and trait diversity (functional richness) compared to larger, less-isolated islands. We also predicted that the trait space of smaller, more isolated islands will be nested within the trait space of larger, less isolated islands, and that the turnover component of trait space dissimilarity will increase with increasing inter-island geographic distance. We also hypothesised that if there is functional redundancy in trait composition — when different species supply the same functions in a community (also known as the ‘insurance’ hypothesis) (McLean *et al*. 2019) — we should observe an asymptotic relationship between trait and species turnover among island pairs; beyond a species-turnover threshold, additional turnover in species would not provide additional turnover in traits under this hypothesis.

## Methods

### Australian islands

We generated spatial data for Australia’s islands and islets and the Australian mainland in QGIS (an open-access, Python-based geographic information system software application: qgis.org; version 3.34.13 ‘Prizren’). We sourced the spatial data for Australia from the GDA2020 digital boundary files from the Australian Bureau of Statistics (Australian Bureau of Statistics 2021). After projecting the shapefile to Australian Albers Equal Area (EPSG:9473), we obtained 9,103 offshore islands and islets within Australian Territory. However, for the purposes of examining both area and isolation effects on species and function richness, we removed some islands according to the following criteria: (*i*) because our objective was to evaluate island biogeographic relationships relative to the Australian mainland source pool, we excluded islands whose nearest major continental landmass was not mainland Australia, including several in the Coral Sea that are closer to New Caledonia, many in Torres Strait/*Zenadth Kes* that are closer to New Guinea, several in the Timor Sea that are closer to Timor, Macquarie Island (and islets) that are closer to New Zealand, Christmas Island and Cocos-Keeling that are closer to Indonesia, several south of Kangaroo Island/*Karta*, Heard, McDonald Islands in the Southern Ocean, the main island of Tasmania/*lutruwitra* as well as all Bass Strait Islands that are closer to Tasmania (e.g., Furneaux group, King Island), and others farther south that are closer to Tasmania than the mainland. This filtering reduces the maximum isolation gradient represented in the data and might contribute to the weak isolation effects observed (see Results). We removed the main island of Tasmania specifically because, with an area of 65,022 km^2^, it is > 11 times larger than the second-largest island in Australia (Melville Island of the Tiwi Islands/*Ratuwati Yinjara*; 5,793 km^2^). After these removals, there were 6,358 islands (69.8%) remaining in the sample (Fig. 1). Most (98.7%; see Results) islands in the dataset were continental shelf islands that were connected to the Sahul landmass during periods of lower sea level.

**Figure 1.**
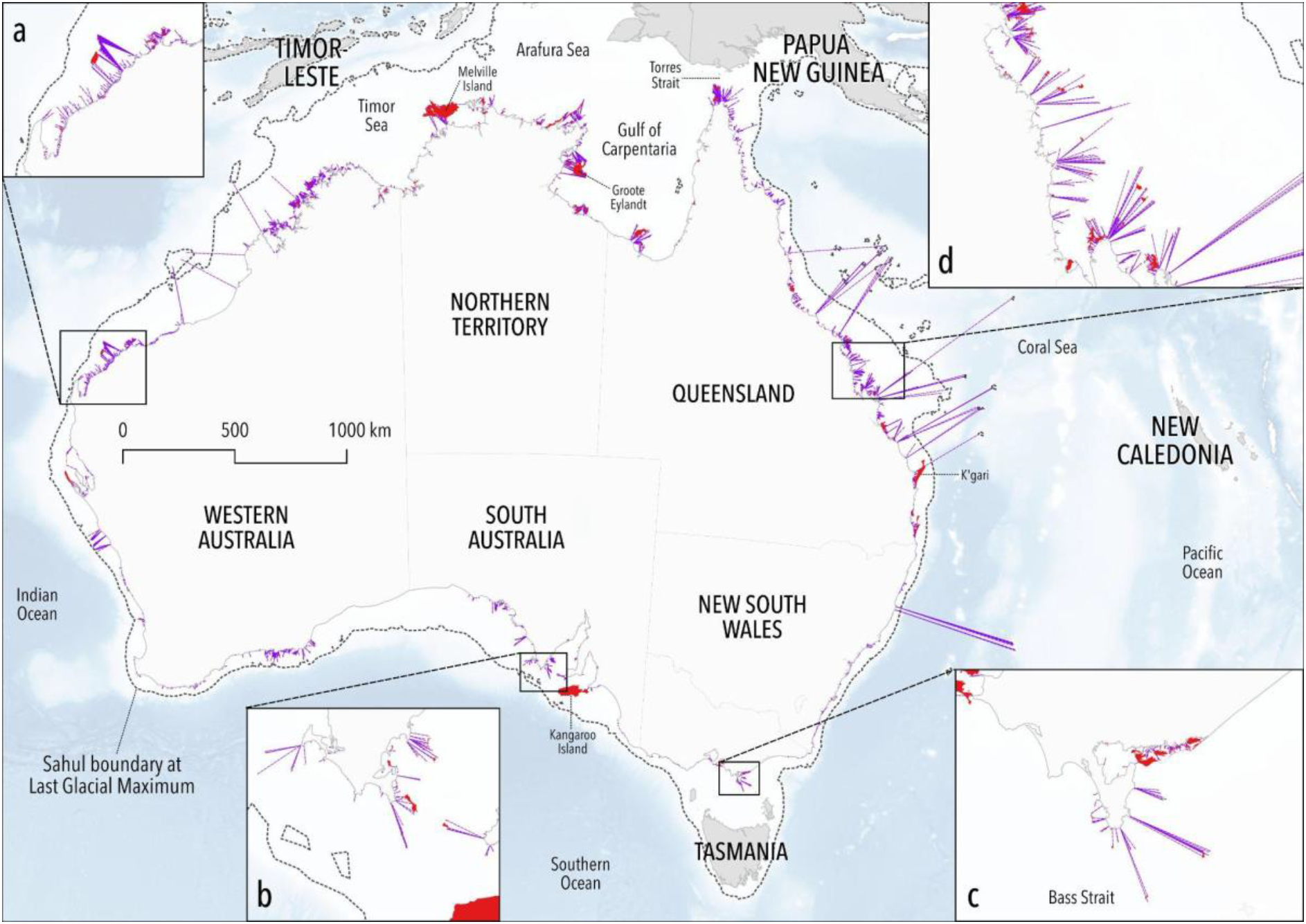
Map of Australia showing the 6,358 islands used in the analysis (red), with their shortest straight-line distance to the mainland shown as dotted purple lines. Insets a–d show zoomed-in views of several different parts of the coastline and their islands. Also shown are major country, state, territory and cadastral features mentioned in the text, as well as the approximate boundary of the supercontinent ‘Sahul’ (Australian mainland + Tasmania + New Guinea joined by dry land) that was exposed during the Last Glacial Maximum approximately 20,000–30,000 years ago (Cadd *et al*. 2021).

We calculated island area in square kilometres (km^2^) using the Australian Albers equal area projection, and the minimum straight-line distance to the Australian mainland using the *shortest line between features* function in QGIS.

### Species data

We used the `galah` package in R (Westgate *et al*. 2025) to access the *Atlas of Living Australia* (ala.org.au) (Belbin *et al*. 2021) database and retrieve occurrence records for mammal, bird, reptile, and amphibian (tetrapod) species located on each island (retrieved 10 July 2025). Given the large size of the occurrence dataset for each taxon (e.g., especially birds: 81,491,093 records for the Australian region), we first removed records occurring within mainland Australia prior to the GIS intersection step to reduce processing time. We then retained island records located within island boundaries using spatial intersection (using the *vector overlay - intersection* tool).

We cleaned the records using the following protocol: (*i*) we removed all non-native introduced species (i.e., introduced/exotic species that did not evolve in Australia; manually checked against species accounts), (*ii*) we removed all marine mammals from the mammal dataset and marine turtles from the reptile dataset, (*iii*) we removed extinct species; (*iv*) we removed all higher-order taxonomic designations that could not specify a unique species (including records only reporting a non-monotypic genus), and (*v*) we reclassified all species with subspecies designation to species-level only.

### Trait data

We obtained trait data for birds and mammals from the *AVONET* (Tobias *et al*. 2022) and *SahulTraits* databases, respectively (*SahulTraits* remains unpublished, but we provide the mammal trait data in the online code and data repository at doi:10.5281/zenodo.21736674). Sufficient trait data for defining trait volumes of Australian reptiles and amphibians are not yet available. In building *SahulTraits*, we collected information on 32 mammal traits from databases and publications, including *COMBINE* (Soria *et al*. 2021), *AnAge* (Tacutu *et al*. 2013), *Strahan’s Mammals of Australia* (Baker & Gynther 2023), and the literature (Squires 1975; Owen-Smith 1988; Boeadi 1990; Macdonald 2001; Woolley 2001; Forsyth *et al*. 2002; Glen & Dickman 2006; Hourigan *et al*. 2006; Cassini *et al*. 2009; Tahir *et al*. 2021; Ablondi *et al*. 2023; Umbrello *et al*. 2023). Completeness of mammal trait data varied from 100% to < 2%. We therefore used multiple imputation — based on measured traits and phylogeny using the `rtrees` R package (Li 2023) — to infer missing data with the `missForest` R package (Stekhoven & Bühlmann 2012). To minimise noise from poorly imputed traits, we restricted our mammal trait-space analysis to traits that had records for > 50% of species (19 traits in total). For the 19 mammal traits we used in the analysis (see below), missingness ranged from 0 to 48.5% (mean = 20.8 ± 19.7% standard deviation). Imputation accuracy of these traits was moderate to high (continuous traits R^2^ = 0.66 to 0.93, categorical traits accuracy = 0.74 to 1.0).

The final set of 19 mammal traits we used to build the trait space included: (*i*) sociality (nominal factor: solitary or gregarious); (*ii*) herbivory (binary factor: yes or no); (*iii*) invertivory (binary factor: yes or no); (*iv*) trophic level (ordinal factor: 1 = herbivore, 2 = omnivore, 3 = carnivore); (*v*) presence of webbed digits (binary factor: yes or no); (*vi*) arboreal locomotion (binary factor: yes or no); (*vii*) terrestrial locomotion (binary factor: yes or no); (*ix*) nocturnally active (binary factor: yes or no); (*x*) crepuscularly active (binary factor: yes or no); (*xi*) log*_e_* body mass (g; numeric); (*xii*) log*_e_* gestation length (days; numeric); (*xiii*) log*_e_* litter size (numeric); (*xiv*) log*_e_*litters/year (numeric); (*xv*) log*_e_* sexual maturity (days; numeric); (*xvi*) residual of the relationship between log*_e_*body mass (g) and log*_e_* total length (mm) (numeric); (*xvii*) residual of the relationship between log*_e_* body mass (g) and log*_e_* longevity (days) (numeric); (*xviii*) residual of the relationship between log*_e_* body mass (g) and log*_e_*weaning age (days) (numeric); and (*xix*) residual of the relationship between log*_e_* body mass (g) and log*_e_* brain mass (g) (numeric).

For birds, we extracted trait data from the *AVONET* database which includes observed and imputed data (Tobias *et al*. 2022). There were 12 bird traits in the final trait dataset: (*i*) habitat structure (ordinal factor: 1 = dense, 2 = semi-open, 3 = open); (*ii*) migration (ordinal factor: 1 = sedentary, 2 = partially migratory, 3 = migratory); (*iii*) primary lifestyle (nominal factor: aerial, terrestrial, insessorial, aquatic); (*iv*) trophic level (nominal factor: herbivore, carnivore, scavenger); (*vi*) hand- wing index = 100*D*_K_/*L*_w_, where *D*_K_ = Kipp’s distance (distance between the tip of the first secondary feather and the tip of the longest primary feather on a folded wing) and *L*_w_ = wing length (distance from the carpal joint to the tip of the longest primary feather) (numeric); (*vii*) log*_e_* body mass (g; numeric); (*viii*) residual of the relationship between log*_e_* body mass (g) and log*_e_* beak length (mm, length from anterior edge of nostrils to tip of beak) (numeric); (*ix*) residual of the relationship between log*_e_* body mass (g) and log*_e_* beak width (mm, width at anterior edge of nostrils) (numeric); (*x*) residual of the relationship between log*_e_*body mass (g) and log*_e_* beak depth (mm, depth at anterior edge of nostrils) (numeric); (*xi*) residual of the relationship between log*_e_* body mass (g) and log*_e_* tarsus length (mm, length of tarsus from posterior notch between tibia and tarsus, to end of last scale of acrotarsium) (numeric); and (*xii*) residual of the relationship between log*_e_* body mass (g) and log*_e_* wing length (mm, length from carpal joint to tip of longest primary on unflattened wing) (numeric).

### Analysis

#### Species-area relationships

We examined species-area relationships for each taxonomic class separately based on the power-law model: *S* = *cA^z^*, where *S* = species richness (total unique species per island), *A* = island area (km^2^), *c* = constant, and *z* = exponent describing the exponential rate of change of *S* with *A*. Although there are many other models considered for the relationship, the power-law form is the most common and is considered robust (Garcia Martin & Goldenfeld 2006). We retained the power model because our goal was to compare slopes across taxa using the most widely used species-area relationship form, while recognising that multi-model SAR frameworks are available for more exhaustive model comparison (Matthews *et al*. 2019b). Although *z* can depend on habitat type, scale, and the taxon under consideration, the most frequently reported values range from 0.2 to 0.4 (Garcia Martin & Goldenfeld 2006; Fattorini *et al*. 2017). To estimate both *c* and *z* for each tetrapod class, we calculated the linear least-squares relationship (function *lm* in R) between log_10_ *S* and (i.e.,). This form estimates both the mean and standard error of the intercept (log_10_ *c*) and slope (*z*). We also tested for possible undersampling of small islands with low recorded species richness by sequentially removing islands with the fewest recorded species and recalculating *z* for each tetrapod taxon. This diagnostic was motivated by the expectation that small or rarely visited islands are more likely to have incomplete occurrence records than larger and more frequently surveyed islands. If low-richness small islands are undersampled, their observed richness will be biased downward, steepening the log-log species-area relationship and inflating the estimated *z* coefficient. Conversely, if low-richness islands are not disproportionately undersampled, removing them should have little systematic effect on *z*.

#### Multivariate models examining combined effects of area and distance

We also employed boosted regression trees (a hybrid machine-learning approach) to quantify the relative importance of area and isolation and to allow for nonlinear responses and interactions between these predictors. Boosted regression trees can model complex, nonlinear relationships and interactions among predictors automatically, without requiring prior specification of these relationships as is necessary in linear or logistic regression (Zhang *et al*. 2019). Boosted regression trees build an ensemble of decision trees iteratively that together improve predictive accuracy (Elith *et al*. 2008). We implemented the boosted regression tree models using the *gbm.step* function from the `dismo` R package (Hijmans *et al*. 2024), setting a Gaussian distribution for the response variable. To optimise the models, we tuned several parameters, including the learning rate (0.0005), tolerance (0.0002), tree complexity (2), and bag fraction (0.75). This balanced robustness of fit with processing speed. We applied cross-validation to assess model performance and to reduce overfitting. All R code and data required to repeat the analyses are available at doi:10.5281/zenodo.21736674.

#### Trait space

For the trait-based analyses, we used the `gawdis` (de Bello *et al*. 2021), `GA` (Scrucca 2013), `FD` (Laliberté *et al*. 2014), and `vegan` (Oksanen *et al*. 2025) packages in R to construct a Gower dissimilarity matrix (Gower & Legendre 1986) that measures the Gower distances between species’ in trait space (i.e., the similarity/dissimilarity in the trait space among species). Gower dissimilarity can handle mixed data types (our trait data included continuous and categorical variables) and can normalise the contribution of variables (Gower & Legendre 1986; Mouillot *et al*. 2021). We then used the `mFD` (Magneville *et al*. 2022) R package to calculate the trait space for each island and quantify trait diversity with the functional richness metric. We constructed separate analyses for the bird and mammal traits. To do this, we first applied principal coordinate analysis to the Gower distances to ordinate species within a lower-dimensional space and calculate functional diversity metrics using `mFD`. We used three principal component axes to retain a consistent trait-space dimensionality across communities while preserving the largest possible sample size. This choice sacrifices some trait-space variance, but avoids excluding species-poor communities and keeps functional diversity metrics comparable. We therefore used these three axes to calculate each community’s functional richness (Mouillot *et al*. 2013).

#### Turnover and nestedness

Species and trait diversity can vary among islands because communities differ in species identities, functional trait composition, or both. We quantified these differences using pairwise dissimilarity among islands and then decomposed dissimilarity into turnover and nestedness components (where total *dissimilarity* = *turnover* + *nestedness*). We first did this decomposition for species composition, and then applied an analogous approach to functional trait space for birds and mammals.

For species composition, we calculated pairwise Jaccard dissimilarity among islands separately for birds, mammals, reptiles, and amphibians using presence-absence matrices. Jaccard dissimilarity measures the proportion of species not shared between two islands relative to the total number of species occurring on either island. We then decomposed total species dissimilarity into turnover and nestedness components following Baselga (2010, 2012). In this framework, turnover represents species replacement between islands resulting from environmental sorting or spatial and historical constraints (Qian *et al*. 2005), whereas nestedness represents differences arising when the species assemblage of a poorer island is largely a subset of that of a richer island (Wright & Reeves 1992; Ulrich & Gotelli 2007; Baselga 2010, 2012). Indeed, nestedness can result from species loss (i.e., absence, not extinction *per se*) (Gaston & Blackburn 2000) or from species gain. We related total species dissimilarity, species turnover, and species nestedness to straight-line distance between island pairs to test whether geographically distant islands support increasingly distinct assemblages.

For functional traits, we applied the same turnover-nestedness logic to the trait spaces of birds and mammals. We first used the principal-coordinate axes derived from the Gower trait-distance matrix to describe the multidimensional trait space occupied by each island community. For each island, we then generated a trait-space hypervolume from the species present on that island. We used pairwise overlap among island hypervolumes to estimate functional dissimilarity, and decomposed this dissimilarity into functional turnover and functional nestedness. Functional turnover indicates replacement of one region of trait space by another between islands, whereas functional nestedness indicates that the trait space of one island is largely contained within that of another. We then related functional turnover and functional nestedness to inter-island distance and to differences in island area. Because turnover and nestedness are complementary components of total dissimilarity, we interpreted their relationships jointly rather than as independent processes. Thus, our central question was whether increasing geographic distance among islands was associated mainly with replacement among assemblages or with increasingly nested subsets of species or trait space.

We therefore expected geographically closer islands to show lower taxonomic and functional turnover, and island pairs differing strongly in richness or area to show stronger nestedness. We then applied the following calculations first to species presence-absence matrices and subsequently to bird and mammal trait hypervolumes. For species composition, we first calculated pairwise Jaccard dissimilarity among islands using presence-absence matrices in the `vegan` R library (function *vegdist* with the ‘jaccard’ option) (Oksanen *et al*. 2025). For a pair of islands *j* and *k*, let *a* be the number of species shared by both islands, *b* the number of species present on island *j* but absent from island *k*, and *c* the number of species present on island *k* but absent from island *j*. Total Jaccard dissimilarity is:

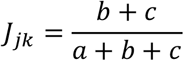

We then decomposed Jaccard’s index into its turnover and nestedness components (Baselga 2012), where the turnover (*τ_jk_*) component is:

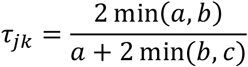

and the nestedness (*ν_jk_*) component is:

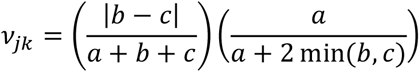

such that:

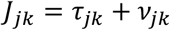

For trait volumes, we took the principal coordinates from the trait-space analyses to build trait-space hypervolumes for each island community using Gaussian kernel density estimation and the `hypervolume` package in R (Blonder *et al*. 2025). We set the requested quantile to 0.95 and estimated the bandwidth for each axis in each island community separately. We then used the mean estimated bandwidth for each axis across the island communities as fixed bandwidths to recalculate hypervolumes (i.e., ensuring the same bandwidths for each island). We used the resulting hypervolumes to quantify distances between community centroids (geometric centre) and overlap in multidimensional trait space. We used the same decomposition logic for functional hypervolumes as for species presence-absence matrices. For all analyses comparing turnover and nestedness, we removed island pairs where turnover or nestedness = 0 or 1, because they skewed the relationship to these extremes, and are unresolved on the logit scale required for linearisation.

## Results

### Species composition

There were 1,661 islands with at least 1 species of any tetrapod taxon recorded in the Atlas of Living Australia (26.1% of the sample of 6,358 islands). These islands had a geometric median area of 0.211 km^2^ (21.1 ha) and ranged from 0.0000974–5794.29 km^2^ (25^th^ percentile = 0.041 km^2^; 75^th^ percentile = 1.50 km^2^; Fig. S1). Geometric median distance to the mainland was 7.8 km for the islands with at least 1 tetrapod species and ranged from 0.5 km to 587.5 km (25^th^ percentile = 10.9 km; 75^th^ percentile = 22.8 km; Fig. S1). Of the 9,103 Australian islands we identified, 8,944 (98.3%) of them were part of the Sahul mainland during the Last Glacial Maximum 20,000–30,000 years ago (and for the 1,661 islands with species data we used in the analyses, 1,640 [98.7%] were once part of the Sahul mainland). So, although these are ‘true’ islands today, they are not permanent oceanic islands subject to colonisation via dispersal. Instead, > 98% of Australia’s islands are continental-shelf islands — fragments of a recently contiguous continental land mass that were isolated as late as 10,000 years ago (Morrison *et al*. 2023). Their tetrapod assemblages can therefore be considered remnants more than colonisers (although substantial colonisation since the Last Glacial Maximum still likely occurred, especially for flighted birds and mammals). Thus, the assemblages on these islands likely reflect species’ historical distributions during periods of lower sea level and within-island dynamics (e.g., persistence and local extinction) rather than mainland-to-island dispersal and colonisation.

When summarised by tetrapod taxonomic class, there were 1,586 islands with ≥ 1 recorded bird species, 608 with ≥ 1 recorded reptile species, 387 islands with ≥ 1 recorded mammal species, and 148 islands with ≥ 1 recorded amphibian species. Geometric median island areas for these respective groups were 0.198 km^2^ (birds), 1.557 km^2^ (reptiles), 3.917 km^2^ (mammals), and 12.912 km^2^ (amphibians) (Fig. S2). These trends approximately correspond with the number of tetrapod species described in the Atlas of Living Australia across all islands (Fig. S3) — i.e., as the number of total species on islands increases per taxonomic group, the number of islands for which there are records increases, and the median size of those islands decreases (Fig. S3). Compared to the number of described species across mainland and island Australia (828 bird species, 985 reptile species, 397 mammal species, and 237 amphibian species) (Dickman 2018), there were 683 bird species recorded on Australian islands in the analysis dataset (84.5% of total bird diversity across Australia), 478 reptile species (48.5% of total reptile diversity across Australia), 177 mammal species (44.5% of total mammal diversity across Australia), and 95 amphibian species (40.1% of total amphibian diversity across Australia) on islands according to the Atlas of Living Australia.

The species-area relationship for each major taxon estimated mean *z* coefficients from 0.217 for amphibians (± 0.027 SE) to 0.309 for reptiles (± 0.013 standard error) (Fig. 2). Although all estimated *z* fell within the expected range of 0.2–0.4 (Garcia Martin & Goldenfeld 2006), the sequential- removal analysis suggested that the lowest-richness islands could exert some influence on slope estimates. If small islands with few Atlas of Living Australia records are undersampled, their observed richness would be biased downward, producing a steeper species-area relationship and an upwardly biased *^Z* (Fig. S4). The observed changes in *^Z* after removing islands with the fewest recorded species are therefore consistent with some undersampling of small or low-richness islands, although the overall slopes remained within expected values (Fig. S4). The broader difference in island-size distributions between islands with Atlas of Living Australia records and all islands further supports this interpretation (Fig. S6).

**Figure 2.**
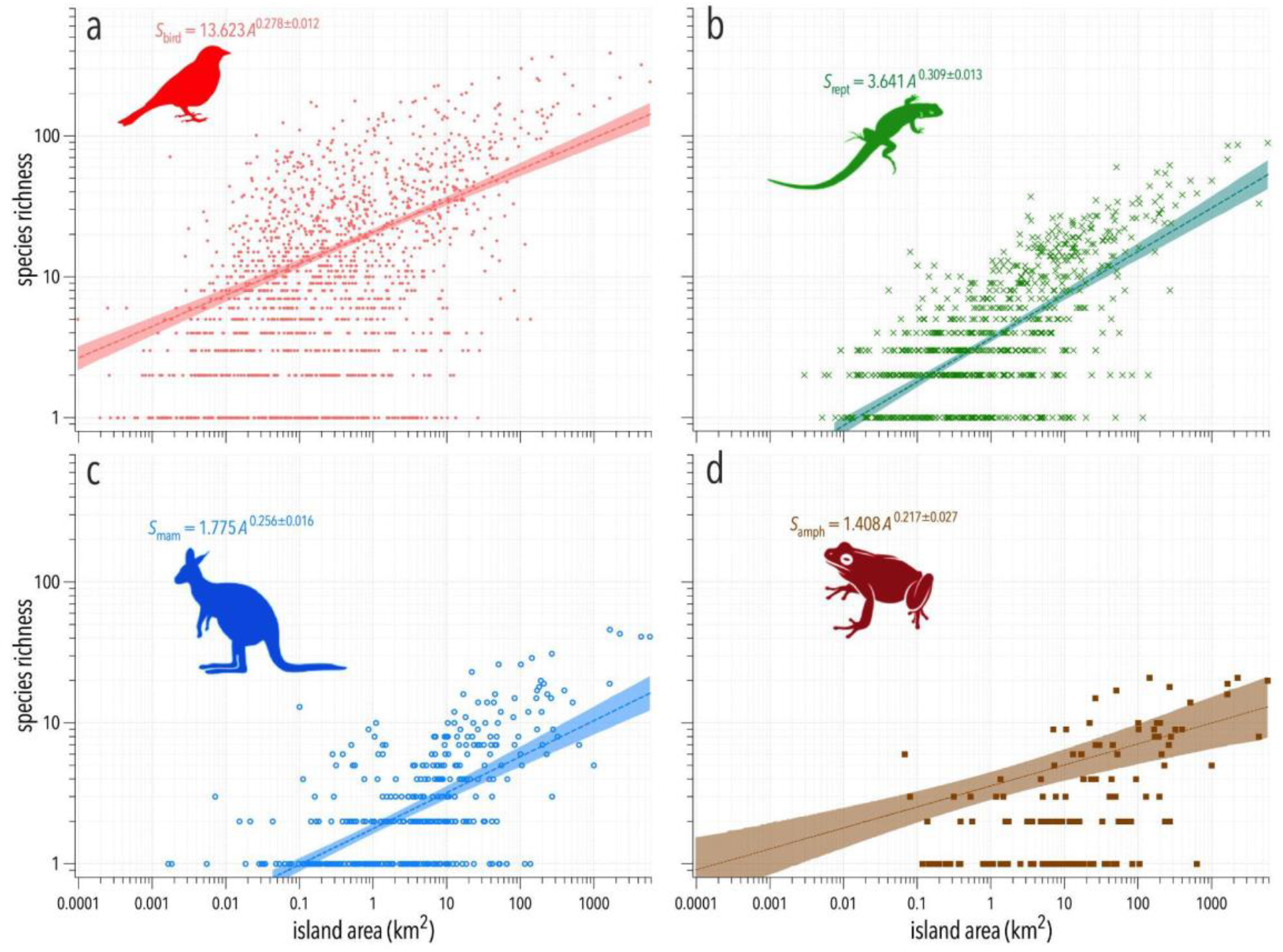
Species richness (*S*, log_10_ scale) as a function of island area (*A*, km^2^; log_10_ scale) for four tetrapod taxa: (a) birds (red), (b) reptiles (green), (c) mammals (blue), and (d) amphibians (brown). R^2^ = 0.26 (birds), 0.47 (reptiles), 0.40 (mammals), 0.31 (amphibians). Each panel also shows the 95% prediction confidence interval for the least-squares line of best fit.

After combining the data from all four taxonomic groups (i.e., total tetrapod species richness), we calculated a *z* = 0.299 ± 0.012 SE (Fig. 3).

**Figure 3.**
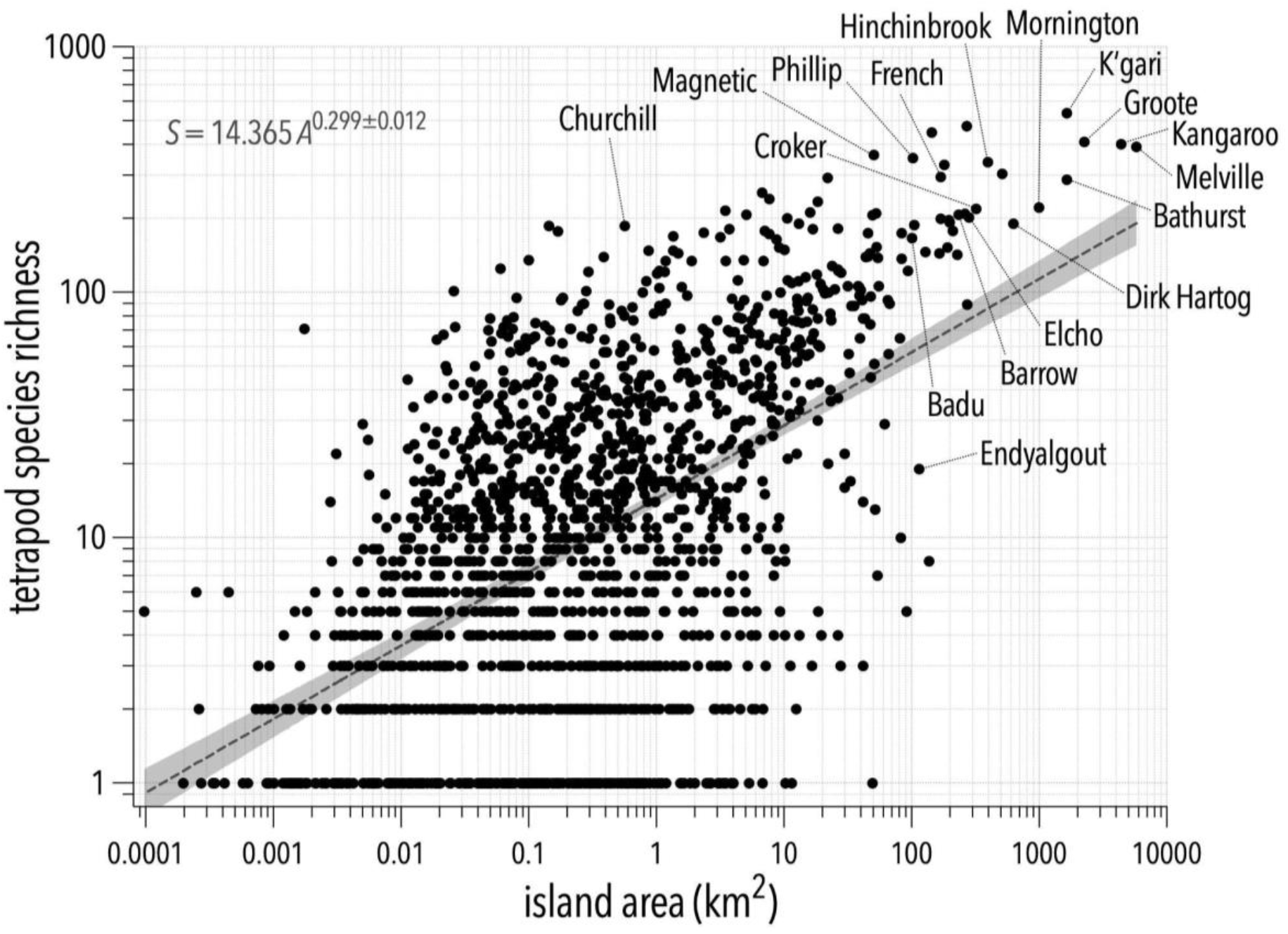
Total tetrapod species richness (*S*; log_10_ scale) as a function of island area (*A*, km^2^; log_10_ scale). R^2^ = 0.28. Named islands: **Badu** Is, Torres Strait, QLD (10.106° S, 142.143° E); **Barrow** Is, WA (20.793° S, 115.394° E); **Bathurst** Is/*Ratuwati Yinjara*, Tiwi, NT (11.645° S, 130.345° E); **Churchill** Is, VIC (38.499° S, 145.337° E); **Croker** Is, NT (11.155° S, 132.543° E); **Dirk Hartog** Is, WA (25.774° S, 113.032° E); **Elcho** Is/*Galiwin’ku*, NT (11.944° S, 135.770° E); **Endyalgout** Is, NT (11.687° S, 132.578° E); **French** Is, VIC (38.355° S, 145.352° E); **Groote** Eylandt/*Ayangkidarrba*, NT (13.994° S, 136.606° E); **Hinchinbrook** Is/*Munamudanamy*, QLD (18.355° S, 146.236° E); **Kangaroo** Is/*Karta*, SA (35.808° S, 137.208° E); **K’gari**, QLD (25.210° S, 153.157° E); **Magnetic** Is/*Yunbenun*, QLD (19.139° S, 146.833° E); **Melville** Is/*Ratuwati Yinjara*, Tiwi, NT (11.609° S, 130.956° E); **Mornington** Is/*Kunhanhaa*, QLD (16.538° S, 139.346° E); **Phillip** Is/*Millowl*, VIC (38.484° S, 145.224° E).

There was no obvious relationship between species richness and isolation (distance to mainland) for any taxon (Fig. S7 and S8). Indeed, the multivariate boosted regression tree analysis including both island area and isolation (minimum straight-line distance to the mainland) as predictors of species richness reveal that island area explains 83.0–98.3% of the explainable variance in species richness across taxa (Fig. S9). Isolation (straight-line distance to the mainland) had almost no (birds: 1.4%), weak negative (mammals, amphibians: 8.7–14.8%), or even a weak positive effect (reptiles: 7.9%) on species richness (Fig. S10).

We first describe spatial patterns in species dissimilarity, turnover, and nestedness for all four tetrapod groups, and then present the equivalent functional-trait analyses for birds and mammals. Overall, dissimilarity in species composition between island pairs increased with increasing distance between islands (Fig. S11). When we decomposed species dissimilarity into its constituent turnover and nestedness components, species turnover increased with geographic distance among islands for all four tetrapod groups, with the strongest relationship observed in mammals and the weakest in birds (Fig. 4). The combined relationship among total dissimilarity, turnover, and nestedness for each taxon is shown in Fig. S12.

**Figure 4.**
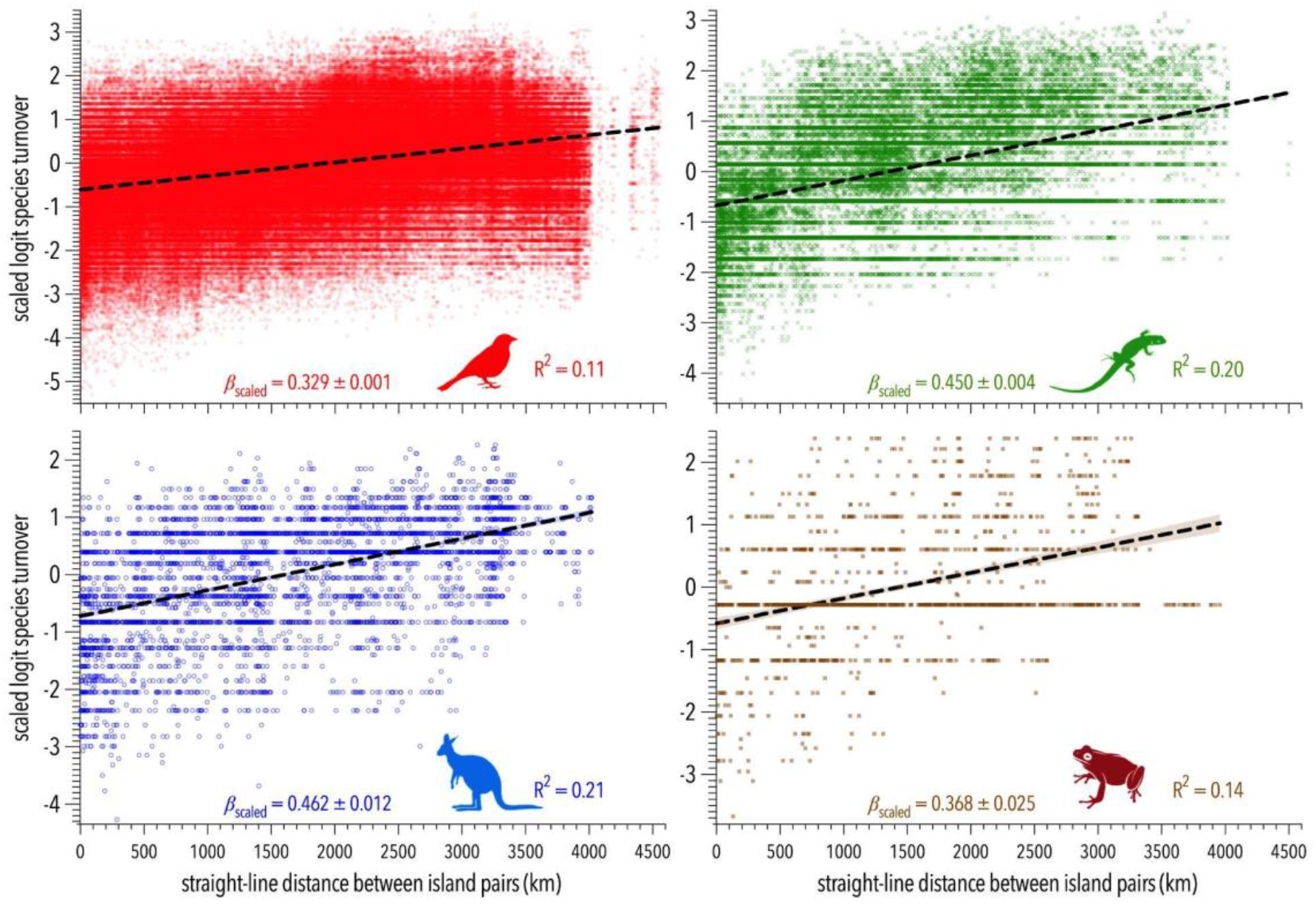
Species turnover (*τ_jk_*; scaled logit) *versus* straight-line distance between island *j*-*k* pairs for the four tetrapod taxa. There were *n* = 514,671, 50,083, 5,140, and 1,415 island pairs for birds, reptiles, mammals, and amphibians, respectively, with sufficient data to estimate species turnover and nestedness (0 < scores < 1).

Species nestedness declined with increasing straight-line distance between island pairs in all four tetrapod groups (Fig. 5), with scaled slopes ranging from -0.149 for amphibians to -0.289 for mammals. Because nestedness and turnover are complementary components of total *β*-diversity, this decline should be interpreted together with the positive distance-turnover relationships. Specifically, as island pairs became more geographically distant, compositional dissimilarity was increasingly explained by species replacement rather than by ordered subset relationships among richer and poorer islands. Thus, distant islands tended to differ because they supported different species, not simply because species-poor islands contained depauperate subsets of species-rich islands.

**Figure 5.**
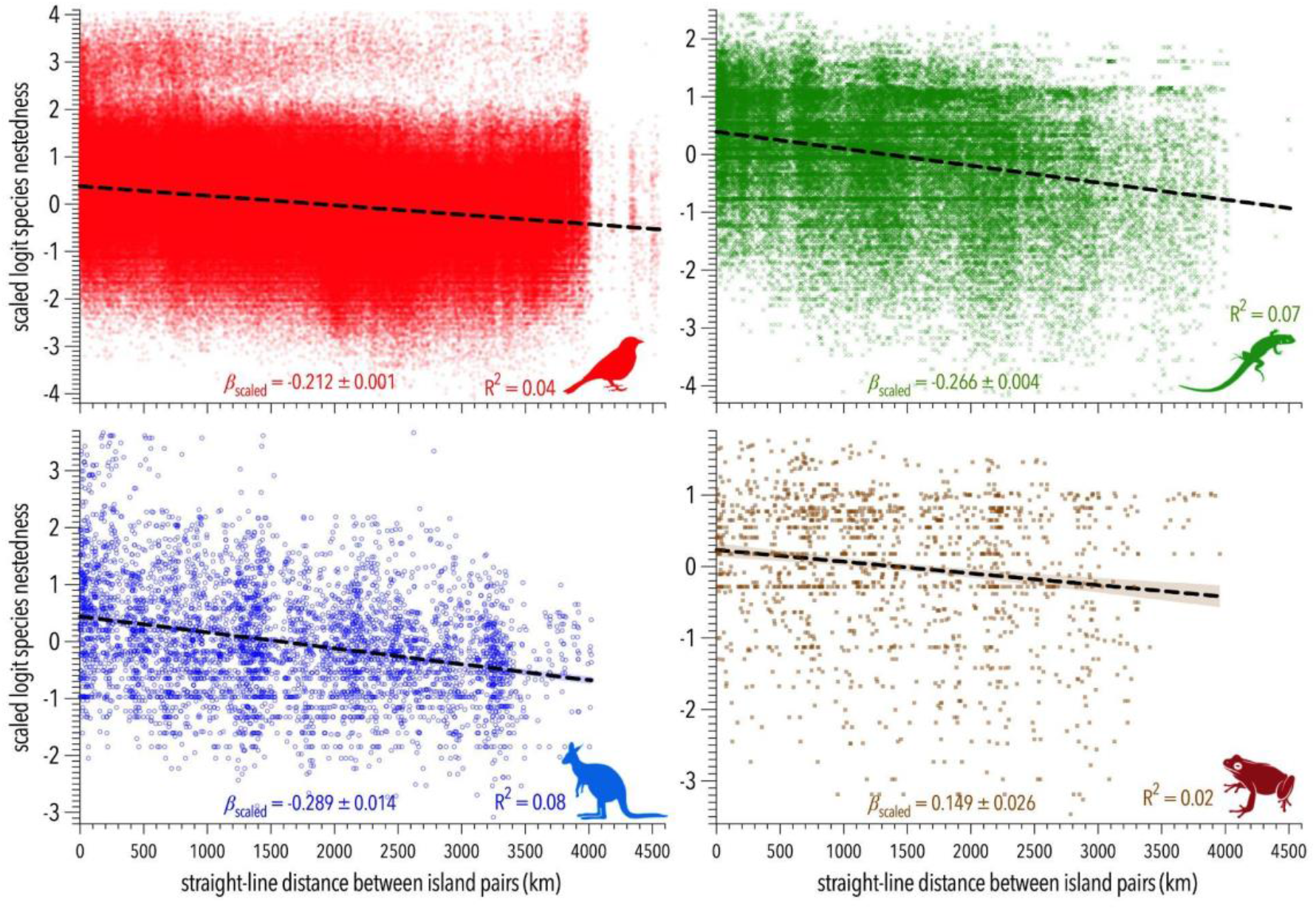
Species nestedness (*ν_jk_*; scaled logit) *versus* straight-line distance between island *j*-*k* pairs for the four tetrapod taxa.

### Trait composition

For mammals, 2 eigenvectors explained 61% of the variation in the functional trait space, 3 explained 70%, and 4 eigenvectors explained 78%. For birds, 2 eigenvectors explained 56% of the variation, 3 explained 67%, and 4 eigenvectors explained 76%. Given we set a lower limit of at least 4 species to construct a trait space per island, we used 3 principal components in the analysis; these 3 components explained 70.4% of the trait variation in mammals and 67.1% in birds, leaving 130 and 1,097 islands for mammals and birds, respectively.

Turnover of traits declined with an increasing absolute difference in island size between pairs for both birds and mammals (Fig. 6), but at a faster rate in mammals (scaled slope *β* = -0.351) than in birds (scaled slope = -0.172). In contrast, trait nestedness increased as the absolute difference in island size increased (Fig. 6), but again at a much faster rate in mammals (scaled slope = 0.285) compared to birds (scaled slope = 0.104).

**Figure 6.**
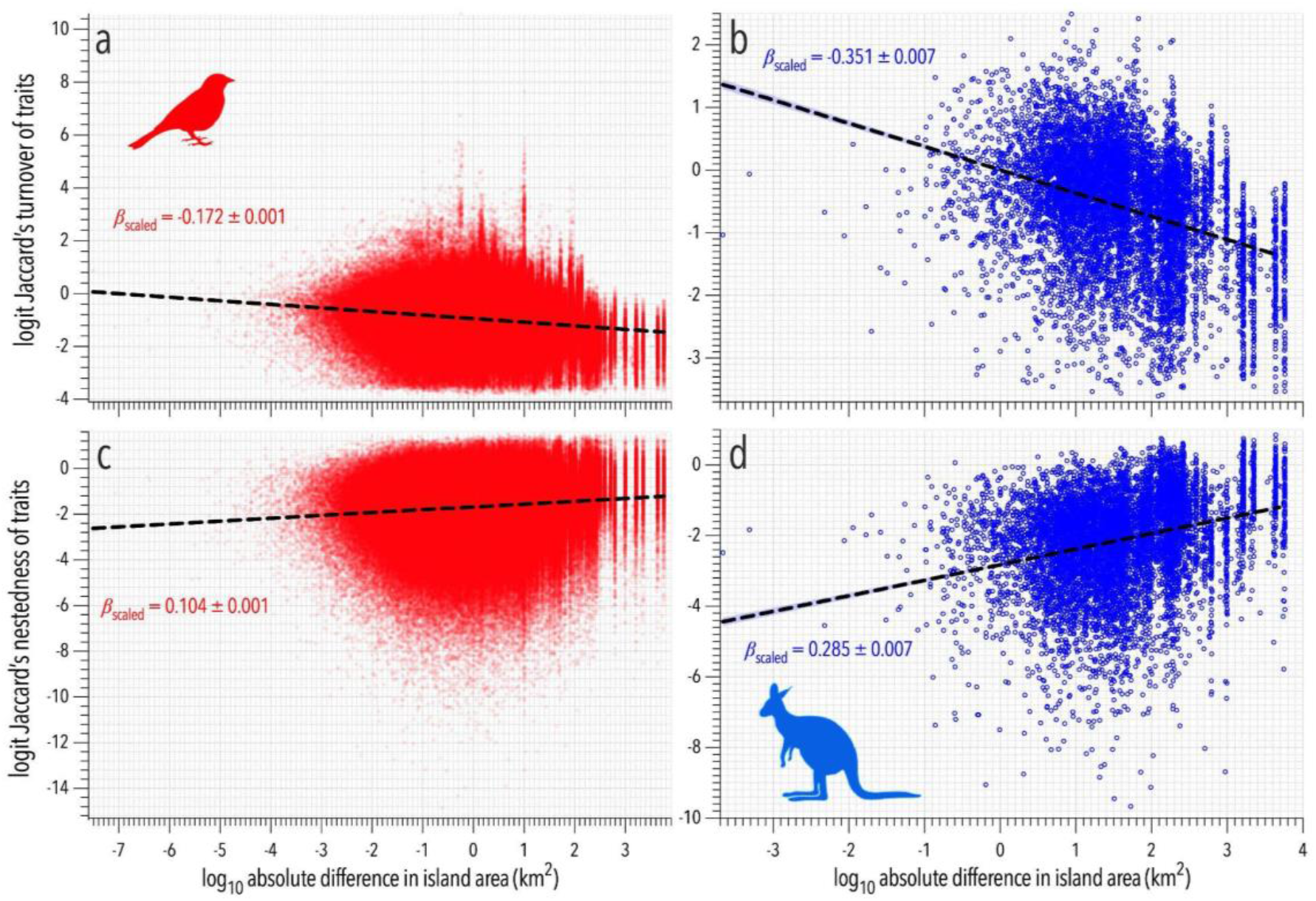
(a, b) Turnover (*τ_jk_*) and (c, d) nestedness (*ν_jk_*) of traits (logit scale) relative to log_10_ absolute difference in area of island *j*-*k* pairs for birds (red) and mammals (blue). Relationship metrics using scaled *y* and *x* variables: (a) bird trait turnover: R^2^ = 0.029, *β* = -0.172 ± 0.001; (b) mammal trait turnover: R^2^ = 0.123, *β* = -0.351 ± 0.007; (c) bird trait nestedness: R^2^ = 0.011, *β* = 0.104 ± 0.001; (d) mammal trait nestedness: R^2^ = 0.081, *β* = 0.285 ± 0.007.

These results demonstrate that (*i*) the trait space of smaller islands tends to be nested in that of larger islands (i.e., islands of similar size are not as nested as those that are different in size), (*ii*) the main differences among similar-sized islands are due to turnover (i.e., replacement, but not loss, of species), and (*iii*) the effects are more pronounced in mammals than in birds.

In contrast, trait turnover increased only modestly with increasing straight-line distance between islands for both birds and mammals, but in this instance at a slightly faster rate for birds compared to mammals (Fig. 7). There was no relationship between trait nestedness and straight-line distance between islands for either birds or mammals (Fig. 7). Trait dissimilarity (turnover + nestedness) increased weakly with distance between islands, and at a slightly higher rate for birds than mammals (Fig. S13).

**Figure 7.**
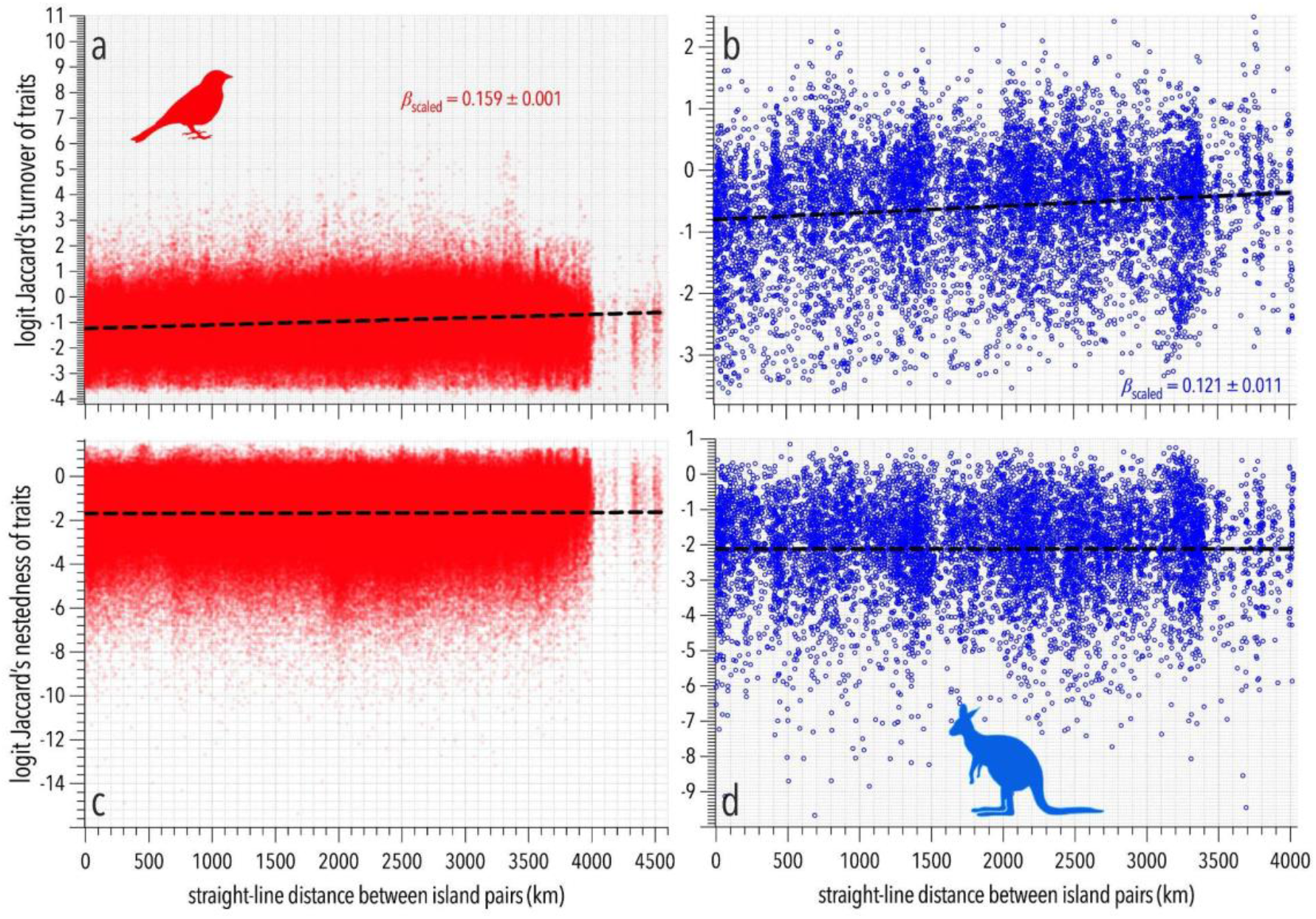
Trait turnover (*τ_jk_*) and nestedness (*ν_jk_*) (logit scale) relative to distance between island pairs *j*-*k*. Relationship metrics using scaled *y* and *x* variables: (a) bird trait turnover: R^2^ = 0.025, *β* = 0.159 ± 0.001; (b) mammal trait turnover: R^2^ = 0.014, *β* = 0.121 ± 0.001; (c) bird trait nestedness: R^2^ = 0.00009, *β* = 0.010 ± 0.001; (d) mammal trait nestedness: R^2^ ≈ 0, *β* = 0.0007 ± 0.011.

### Trait versus species composition

Functional richness for birds and mammals increased with increasing island area, but the relationship was slightly weaker than for species richness (Fig. 8). When scaled to compare the *z* coefficients for the species-area and functional richness-area relationships (using the logit for the latter because of the measure’s constraint of 0–1), functional richness increased at approximately the same rate (scaled *z* = 0.555 ± 0.086) as species richness (scaled *z* = 0.655 ± 0.087) with island area for mammals (i.e., scaled *z* coefficients indistinguishable considering parameter uncertainty; Fig. 8, top panel). For birds, functional richness increased at a slightly lower rate (scaled *z* = 0.405 ± 0.024) than species richness (scaled *z* = 0.481 ± 0.023) with island area, even after scaling and accounting for the truncated distribution of functional richness ∼ 1 (Fig. 8).

**Figure 8.**
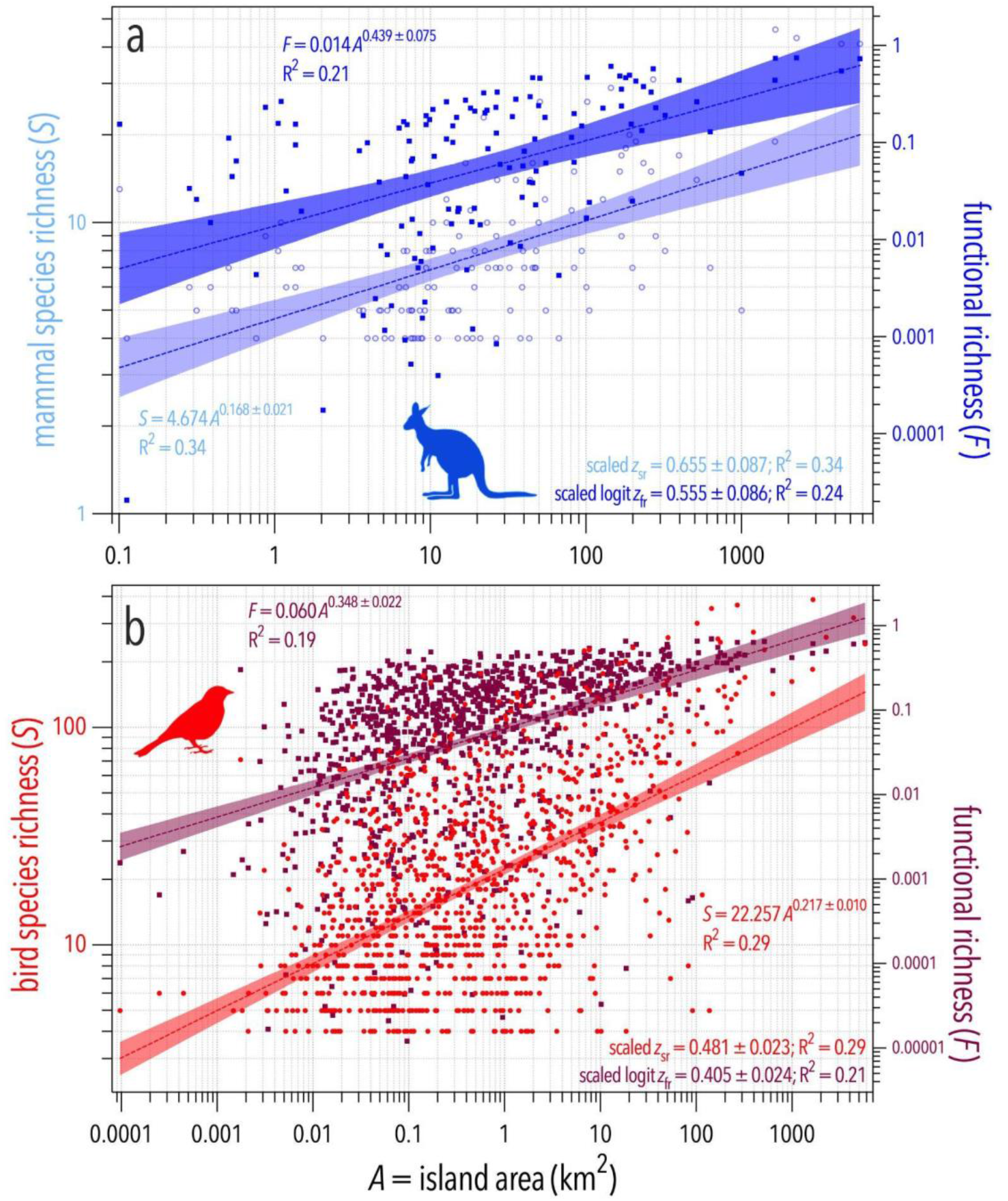
Species (*S*; left *y* axis; log_10_ scale) and functional richness (*F*; right *y* axis; log_10_ scale) plotted against island area (km^2^; log_10_ scale) for mammals (a) and birds (b). Only islands including at least 4 species of the relevant taxon included. Shown are the estimated power-law species-area (blue = mammals; red = birds) and function diversity-area (purple) equations in each panel (*c* scaling and *z* coefficients) and the goodness of fit (R^2^) for each log-log relationship. Also shown are the scaled ([*x* - *̅x*]/*σ_x_*) *z* coefficients for the species- (*z*_sr_) and functional (*z*_fr_) richness-area relationships (the latter is first logit-transformed because of its 0–1 range) so that they can be compared directly.

Functional richness increased strongly with species richness for both birds and mammals (birds: R^2^ = 0.688, P < 0.001; Akaike’s information criterion corrected for small samples [AIC*_c_*] evidence ratio [ER = AIC*_c_* weight of the slope-intercept model ÷ AIC*_c_*weight of the intercept-only model] = 5.8×10^275^; mammals: R^2^ = 0.570, P < 0.001; ER = 2.3×10^22^; Fig. S14). However, island area explained no residual variation in functional richness after accounting for species richness in either birds (type I error estimate [P] = 0.320) or mammals (P = 0.735). Likewise, island isolation explained no additional variation in functional richness once species richness was included in the models (birds: P = 0.543; mammals: P = 0.568). These results indicate that relationships between island geography and functional richness are largely mediated through species accumulation rather than through direct effects on functional trait space.

As for species richness, there was no relationship between functional richness and minimum straight-line distance to the mainland for either birds or mammals (Fig. S15).

Indeed, there was evidence for a non-linear (quadratic) relationship between trait turnover and species dissimilarity (Fig. S16). When we decomposed species dissimilarity into turnover and nestedness components, boosted regression tree fits revealed asymptotic behaviour of trait turnover relative to species turnover for both birds and mammals (Fig. 9; see also Fig. S16 for relative linear and quadratic fits). However, trait nestedness increased near-linearly with species nestedness for both birds and mammals (Fig. S17).

**Figure 9.**
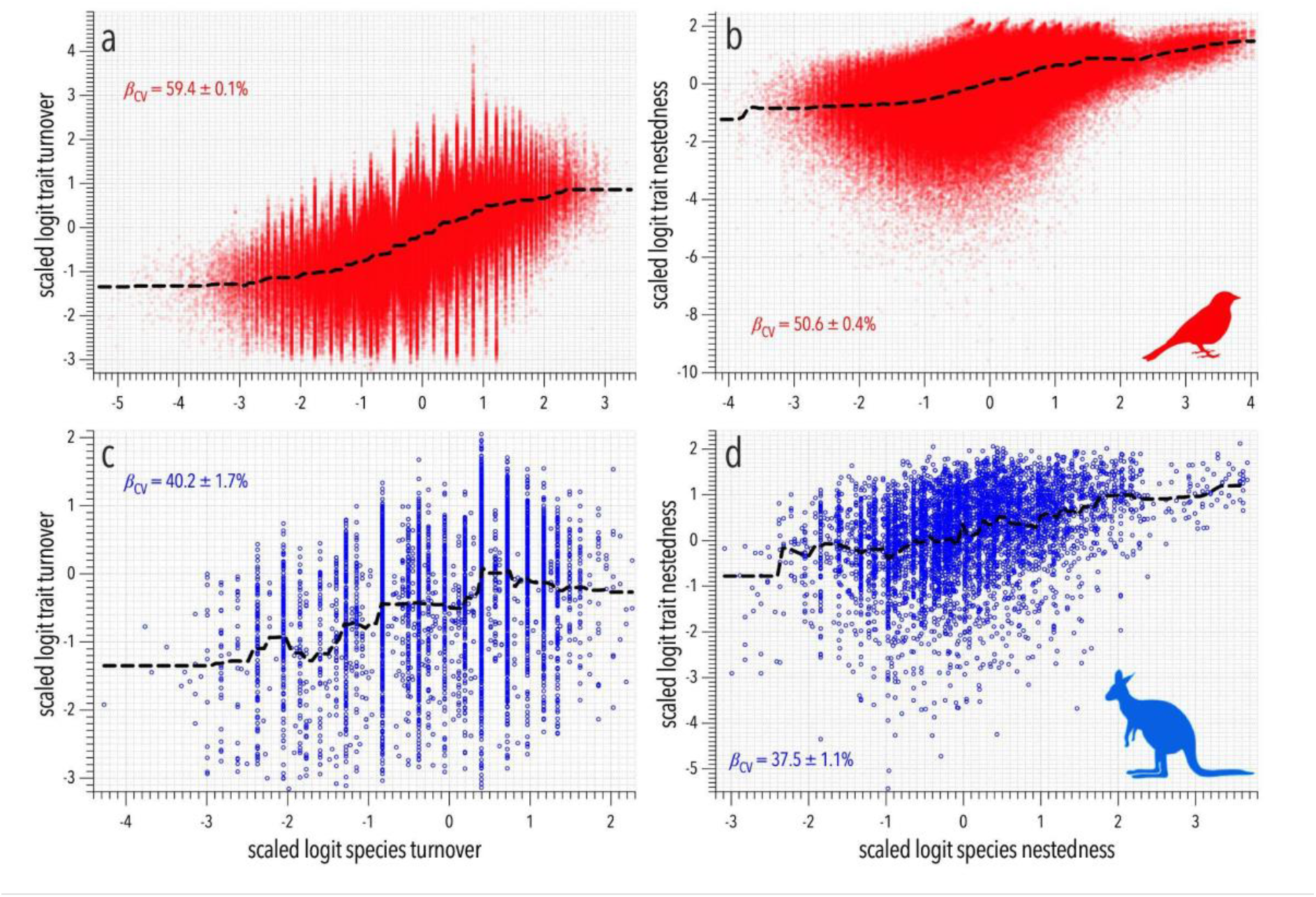
Trait turnover and nestedness (scaled logit) relative to species turnover and nestedness for both birds (red) and mammals (blue). Boosted regression tree models fitted to each relationship, with the coefficient of variation (*β*_CV_) (goodness of fit) shown for each. Dotted line is the fitted deterministic partial relationship predicted from the boosted regression tree.

## Discussion

Island biogeography theory predicts that diversity patterns emerge from the interacting effects of island area and isolation on extinction and colonisation processes (MacArthur & Wilson 1967; Whittaker *et al*. 2017; Matthews & Triantis 2021). Indeed, global mammal-island analyses show that past isolation can be important for present-day mammal diversity, particularly for non-volant species (Barreto *et al*. 2021). However, the Australian islands we examined differ from the oceanic archipelagos that motivated much of classical island biogeography theory. Most (> 98%) islands in our dataset were connected to mainland Australia during periods of lower sea level associated with Quaternary glacial cycles, indicating that contemporary vertebrate assemblages are largely products of relatively recent fragmentation of continental biotas rather than long-term colonisation dynamics (Cadd *et al*. 2021; Morrison *et al*. 2023). In contrast to oceanic island floras where colonisation and speciation can generate substantial trait-space expansion (Barajas Barbosa *et al*. 2023), Australian continental-shelf tetrapod assemblages appear to reflect species persistence and local extinction more strongly. This interpretation is also consistent with global analyses showing that local and intra-archipelago processes can obscure expected isolation effects in island species-area relationships (Matthews *et al*. 2019a). Interpreting Australian islands within this continental-shelf context explains why the relative importance of area and isolation differed from many expectations derived from remote oceanic island systems (Whittaker *et al*. 2017; Matthews & Triantis 2021).

Consistent with this interpretation, island area emerged as the dominant predictor of vertebrate richness across all four taxonomic groups. Species-area relationships were positive and remarkably consistent among birds, mammals, reptiles, and amphibians, and area was also the most influential predictor in the boosted regression tree analyses. These findings accord closely with classical species-area theory predicting greater richness on larger islands because they support larger populations, greater environmental heterogeneity, and lower extinction probabilities (MacArthur & Wilson 1967; Lomolino 2001). Larger islands also provide a greater diversity of habitats and ecological opportunities, allowing species with different environmental requirements to coexist (Triantis *et al*. 2003; Fattorini *et al*. 2017). The consistency of area effects across all vertebrate groups suggests that common ecological processes underpin diversity patterns throughout Australian island systems.

We also found a close correspondence between taxonomic and functional diversity patterns. Functional richness increased strongly with island area and displayed similarly weak responses to isolation, indicating that the same geographical factors that shape species richness also influence the diversity of ecological strategies represented within island communities. Functional diversity has increasingly been recognised as an important complement to taxonomic diversity because it captures variation in ecological roles rather than just the number of species present (Violle *et al*. 2007; Mammola *et al*. 2021). However, additional analyses revealed that the relationships between geography and functional richness were mostly mediated through species richness. Functional richness was strongly related to species richness in both birds and mammals, with species richness explaining ∼ 69% and 57% of variation in functional richness, respectively. In contrast, neither island area nor isolation explained additional variation in functional richness after accounting for species richness. This result is consistent with the global avian island patterns (Matthews *et al*. 2023) showing that increasing species richness with area is often the primary driver of apparent functional and phylogenetic diversity-area relationships. Thus, geographic effects on functional diversity appear to operate largely through the accumulation and persistence of species rather than through independent expansion or contraction of functional trait space. These findings support emerging trait-based extensions of island biogeography theory and suggest that the processes shaping species richness also govern the breadth of ecological functions represented within island assemblages (Schrader *et al*. 2021; Schrader *et al*. 2023).

Patterns of *β*-diversity further illuminate the processes structuring Australian island communities. Species turnover increased with geographic distance among islands, indicating that more distant islands tend to support increasingly distinct species assemblages. We observed similar patterns for functional turnover, suggesting that the replacement of species among islands is accompanied by corresponding shifts in ecological trait composition. Partitioning *β*-diversity into turnover and nestedness components provides important insights into the mechanisms generating compositional differences among communities (Baselga 2010, 2012). We found that turnover generally contributed more strongly than nestedness, implying that differences among island assemblages are driven primarily by species replacement rather than by ordered loss of diversity. This result is consistent with large-scale biogeographic studies demonstrating that environmental variation and geographic separation frequently generate turnover-dominated patterns across landscapes (Qian *et al*. 2005). Similarly, the relatively weak nestedness signal suggests that species-poor islands are not merely subsets of richer islands, but often contain distinct assemblages and combinations of ecological traits (Wright & Reeves 1992; Ulrich & Gotelli 2007).

The strong concordance between taxonomic and functional diversity patterns has important implications for biodiversity conservation. Conservation priorities are often identified based on species richness, yet species numbers alone do not necessarily reflect the range of ecological functions maintained within communities (Violle *et al*. 2007; Mammola *et al*. 2021). For Australian islands, our results suggest that areas supporting high species richness also tend to support broader functional trait diversity. Because functional richness was largely explained by species richness, conservation actions that maintain species-rich island assemblages are also likely to preserve a substantial proportion of ecological functions. Consequently, the protection of larger islands should conserve both taxonomic diversity and functional diversity simultaneously, even though island area does not appear to exert an independent influence on functional richness once species richness is considered.

Several limitations with the data should be acknowledged. We were obliged to restrict functional analyses to birds and mammals because comparable trait datasets are currently unavailable for most Australian reptiles and amphibians. Consequently, whether the correspondence between taxonomic and functional diversity extends across all vertebrate groups remains uncertain. In addition, the predominance of continental shelf islands in our dataset means that our conclusions might not be directly transferable to highly isolated oceanic archipelagos where colonisation processes play a much larger role in community assembly. Contemporary island communities could also reflect anthropogenic impacts, including habitat modification, introduced predators, and historical extirpations. This is relevant for Australian islands because introduced mammals such as cats, foxes, and rodents are major threats to native island biodiversity (McCreless *et al*. 2016; Spatz *et al*. 2017; Holmes *et al*. 2019). Such impacts could depress observed species richness, especially for mammals and birds, and thereby distort island area-richness and area-functional richness relationships if local extirpations have occurred disproportionately on islands with stronger impacts from human activity, invasive-species establishment, or predator-management intensity. Because we could not include island-specific histories of invasive predators, eradication, habitat modification, or native species loss, our richness estimates should be interpreted as contemporary realised native assemblages rather than as pre-disturbance island faunas. Future studies incorporating explicit data on introduced predators, eradication histories, habitat disturbance, and historical species records would help distinguish biogeographic effects of island area and isolation from human-mediated losses.

Overall, our results demonstrate that taxonomic and functional diversity on Australian islands respond to geography in remarkably similar ways. The predominance of continental shelf islands means that island area, rather than isolation, is the principal determinant of vertebrate diversity. Importantly, the influence of island geography on functional richness appears to be almost entirely mediated through species richness. For both birds and mammals, island area and isolation ceased to explain functional richness after accounting for species richness, indicating that geographic processes shape ecological function primarily through their effects on species accumulation and persistence. These findings highlight the enduring influence of Quaternary geological history on contemporary biodiversity patterns and reinforce the close linkage between taxonomic and functional dimensions of diversity across Australian island systems. More broadly, they support the emerging view that island-biogeographic processes operate not only on the number of species present, but also on the structure of the functional trait space those species collectively occupy (Whittaker *et al*. 2017; Schrader *et al*. 2021; Schrader *et al*. 2023).

## Acknowledgements

Funding for developing the *SahulTraits* mammal trait database obtained from the Australian Research Council Centre of Excellence for Australian Biodiversity and Heritage (2017–2025; CE170100015; epicaustralia.org.au). We acknowledge the sovereign Traditional Owners and custodians (First Nations) of the unceded lands, seas and skies where we live and work, including Kaurna in Tarndanya/Adelaide (C.J.A.B., J.L., C.M., A.N.), Peramangk in Bukatila/Mount Lofty Ranges (C.J.A.B.), Dharawal in Woolungah/Wollongong (F.S.), Dharawal/Sydney (F.S., C.R.D.), and Turrbul in Meeanjin/Brisbane (A.E.R., H.P.P.).

## Data and code availability

All code and data necessary to repeat the analyses available at doi:10.5281/zenodo.21736674.

## Supplementary information

**Figure S1.**
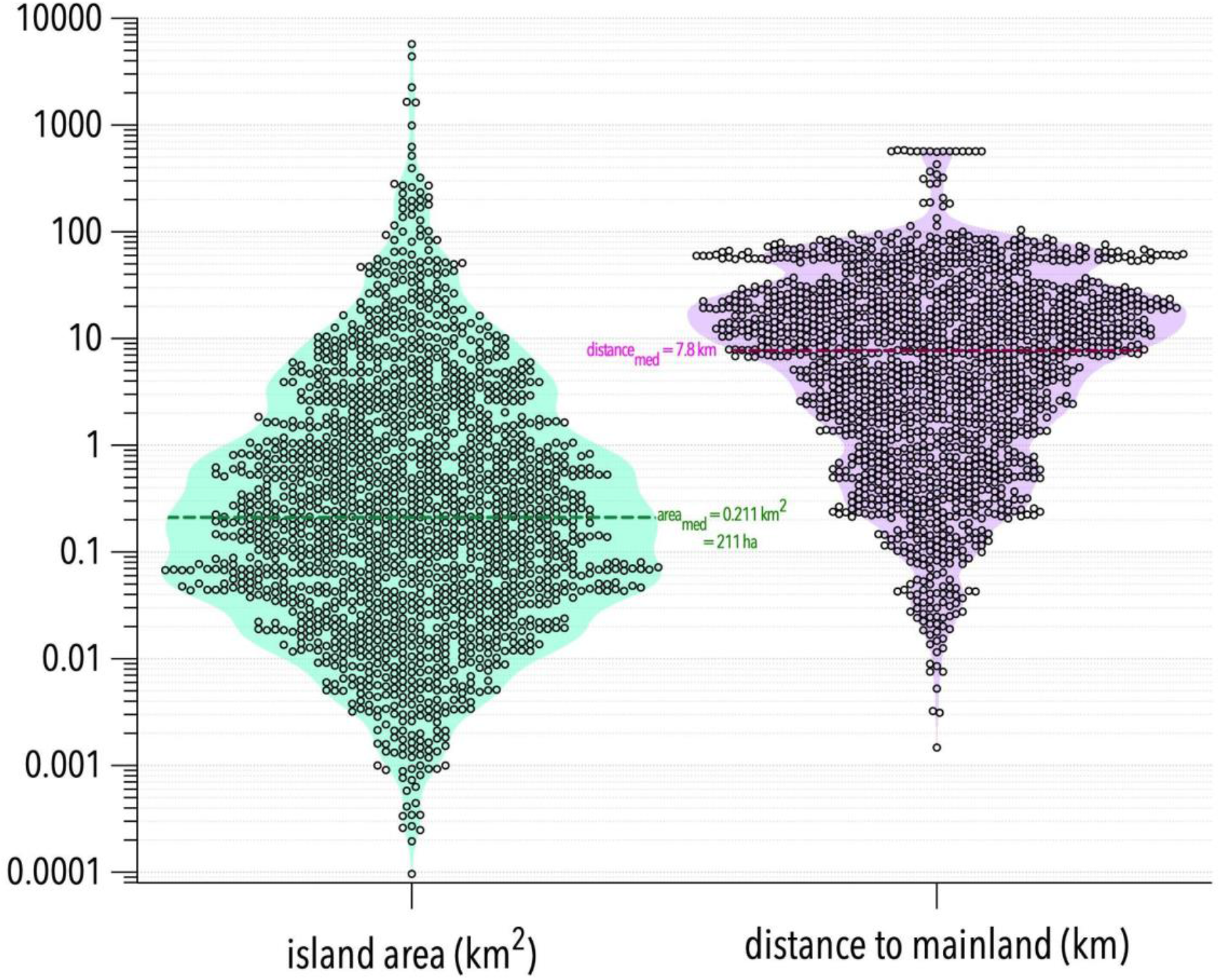
Violin plots of total area (green, km^2^; log_10_ scale) and minimum straight-line distance to mainland (purple, km; log_10_ scale) for 6358 islands around Australia. Shown for each distribution is the geometric median (area_med_ and distance_med_).

**Figure S2.**
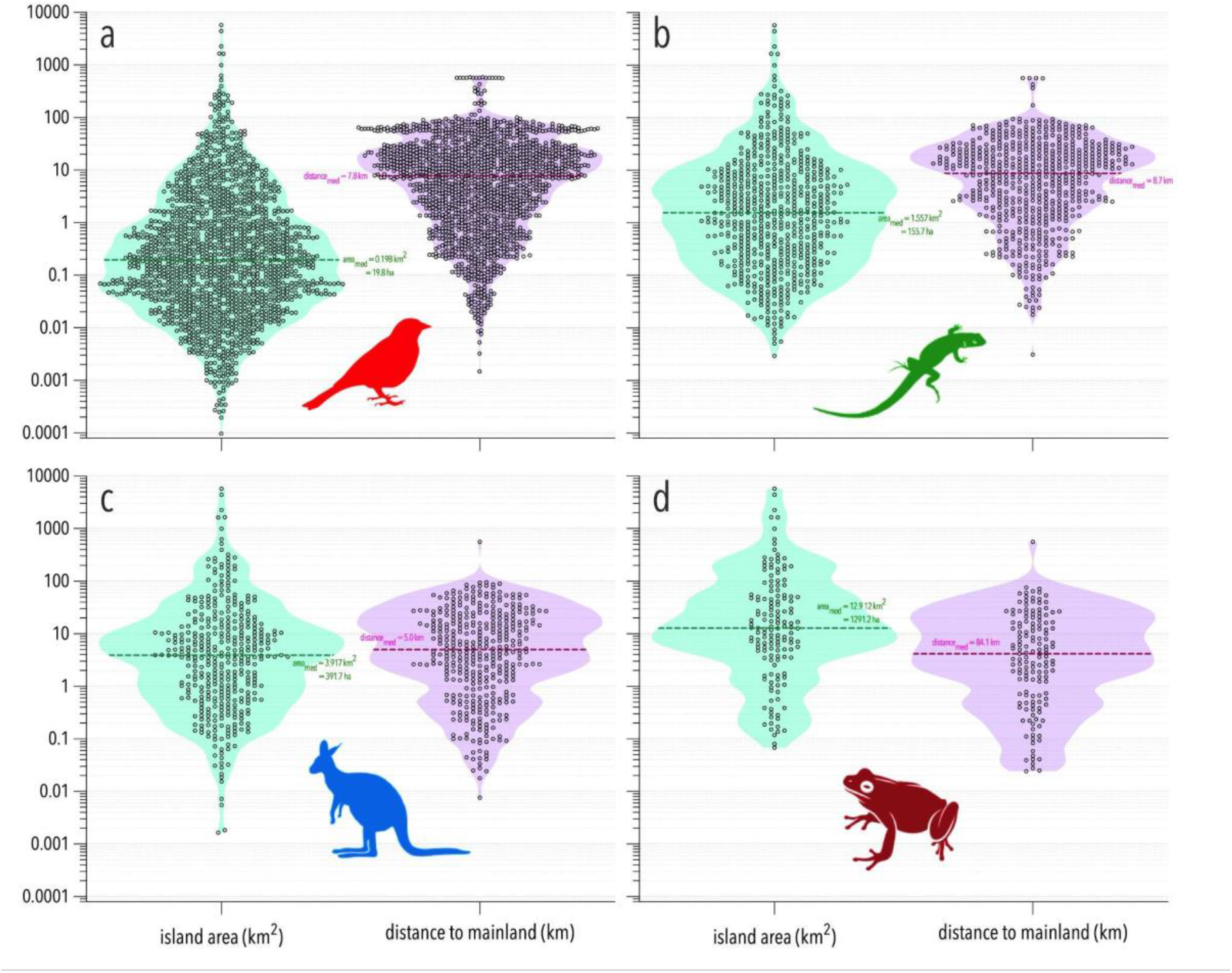
Violin plots of total area (green, km^2^; log_10_ scale) and minimum straight-line distance to mainland (purple, km; log_10_ scale) for islands around Australia with (a) ≥ 1 bird species (*n* = 1,586 islands), (b) ≥ 1 reptile species (*n* = 608 islands), (c) ≥ 1 mammal species (*n* = 387 islands), and (d) ≥ 1 amphibian species (*n* = 148 islands). Shown for each distribution for each species is the geometric median (area_med_ and distance_med_).

**Figure S3.**
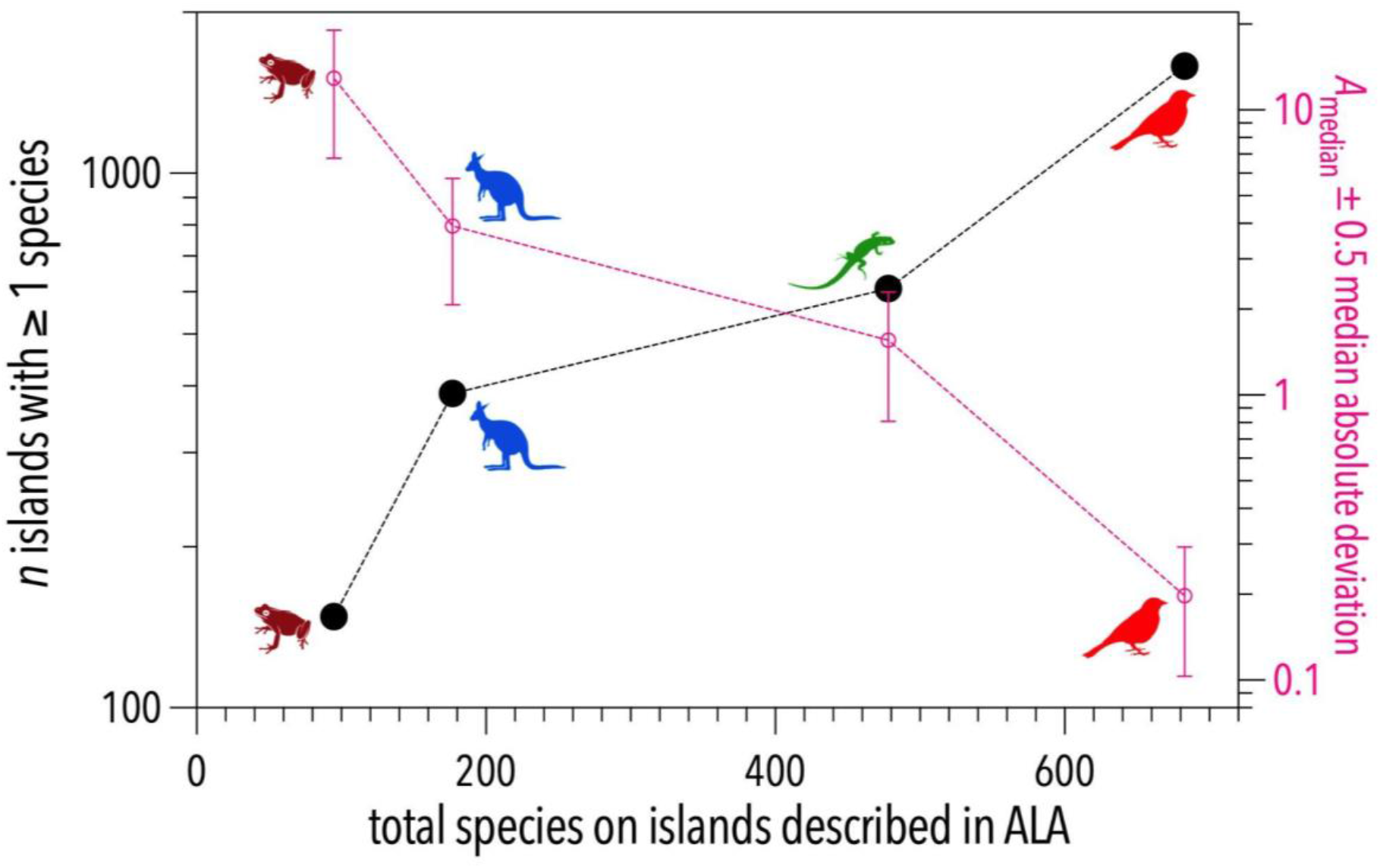
Relationship between number of islands (left *y* axis; log_10_ scale) with at least 1 species per major tetrapod taxon (birds, reptiles, mammals, amphibians) and the number of species in the Atlas of Living (ALA) on the islands in our sample Australia (*x* axis), and median (± 0.5 median absolute deviation) island area (right *y* axis). Compared to the number of described species across Australia (828 bird species, 985 reptile species, 397 mammal species, and 237 amphibian species; Dickman 2018), there were 683 bird species (84.5% of total bird diversity across Australia), 478 reptile species (48.5% of total reptile diversity across Australia), 177 mammal species (44.5% of total mammal diversity across Australia), and 95 amphibian species (40.1% of total amphibian diversity across Australia) on islands according to the Atlas of Living Australia.

**Figure S4.**
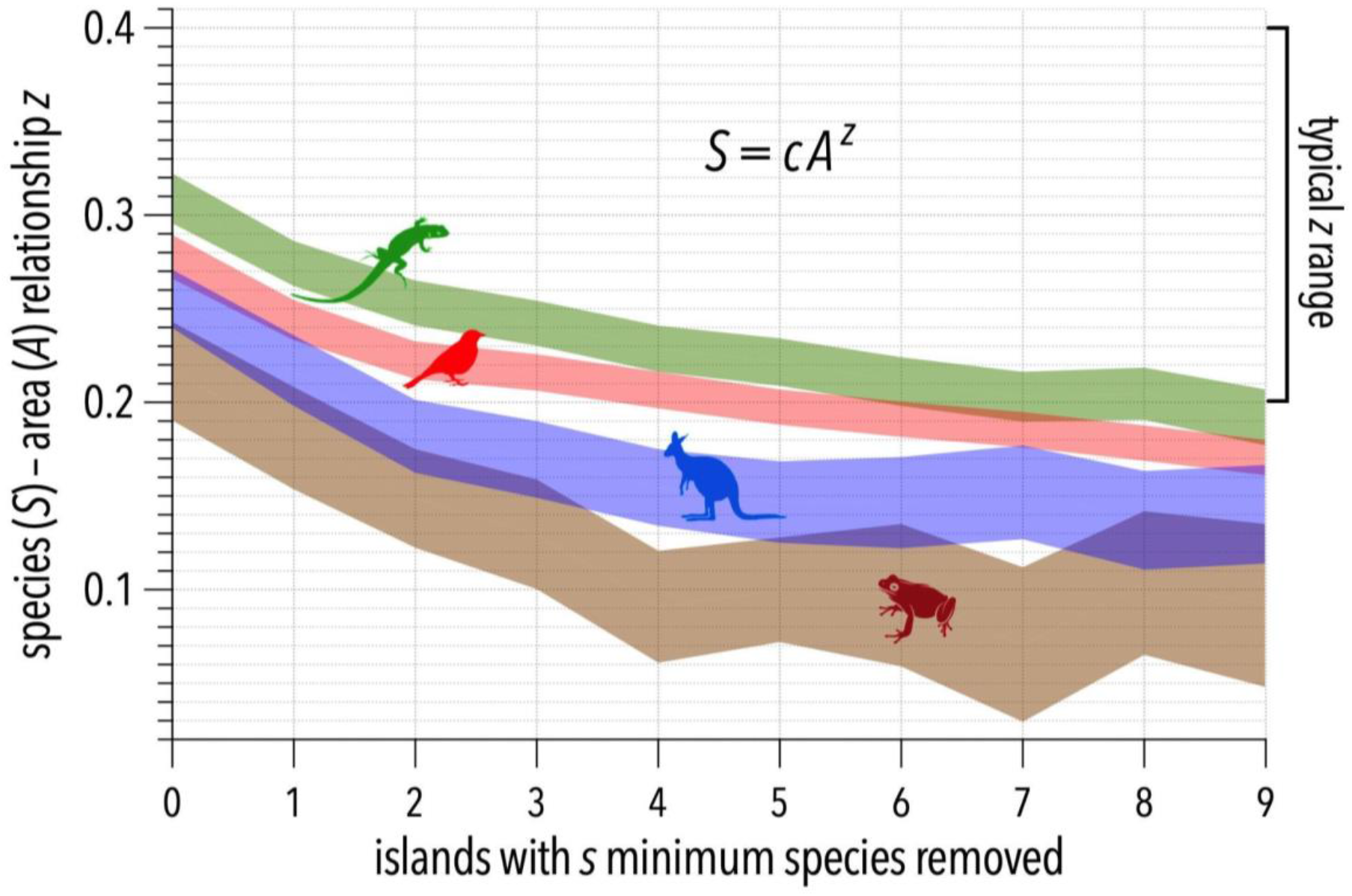
Change in the estimated value of *z* exponent of the power-law species-area relationship *S* = *cA^z^* (where *S* = species richness, *c* = a constant, and *A* = island area) as islands with the lowest number of species recorded in the Atlas of Living Australia are removed sequentially.

**Figure S5.**
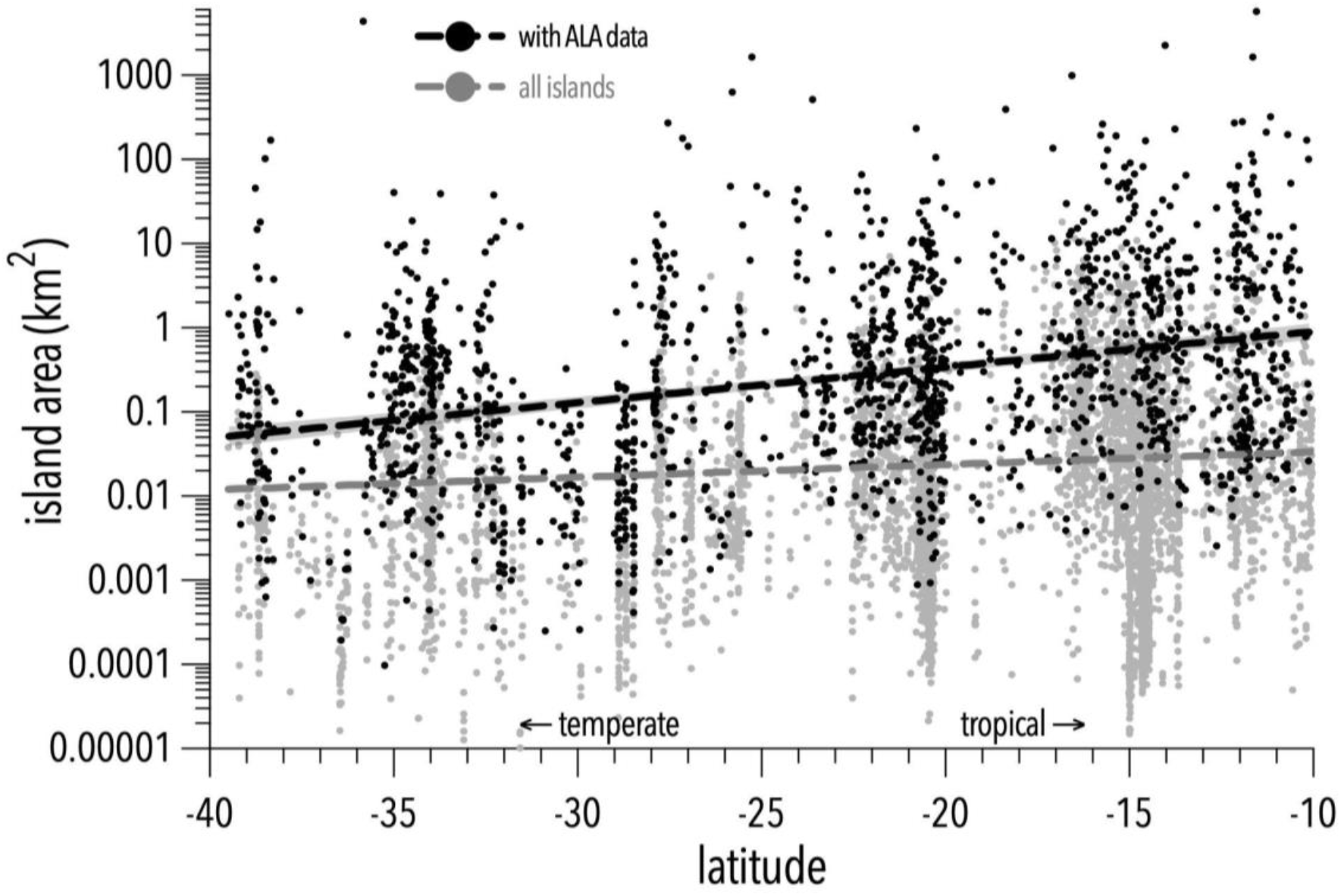
Island area (km^2^; log_10_ scale) versus latitude. Islands with Atlas of Living Australia (ALA) data at 10 °S latitude are 17.3× larger than those at 40 °S latitude on average (R^2^ = 0.096; evidence ratio ≈ ∞); for the all-islands sample, those at 10 °S latitude are 2.8× larger than those at 40 °S latitude on average (R^2^ = 0.009; evidence ratio ≈ ∞).

**Figure S6.**
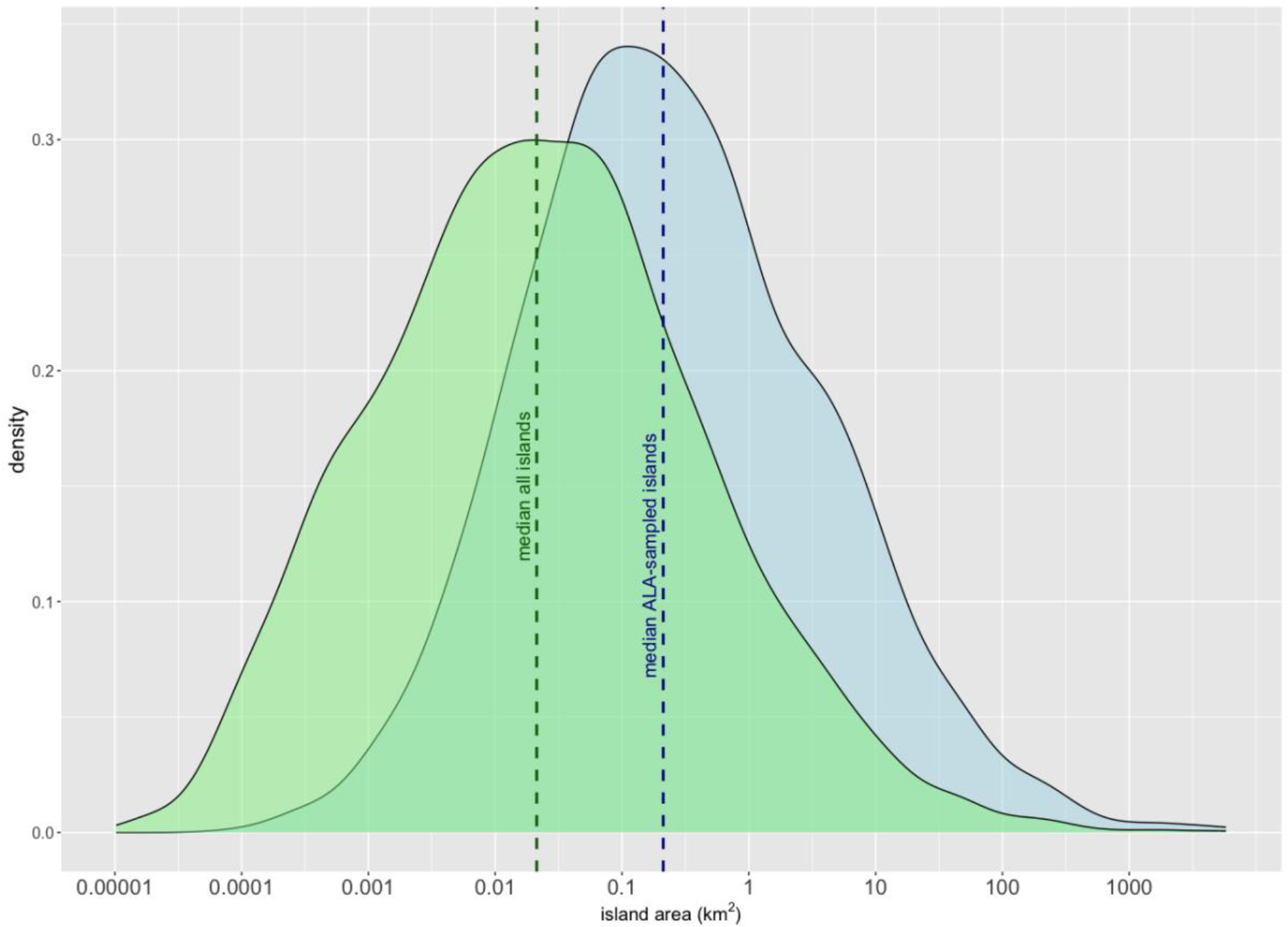
Probability densities of island area (km^2^; log_10_ scale). Islands with Atlas of Living Australia (ALA) data (blue distribution) are a median of one order of magnitude larger than the total sample of islands we used in the analysis (green distribution).

**Figure S7.**
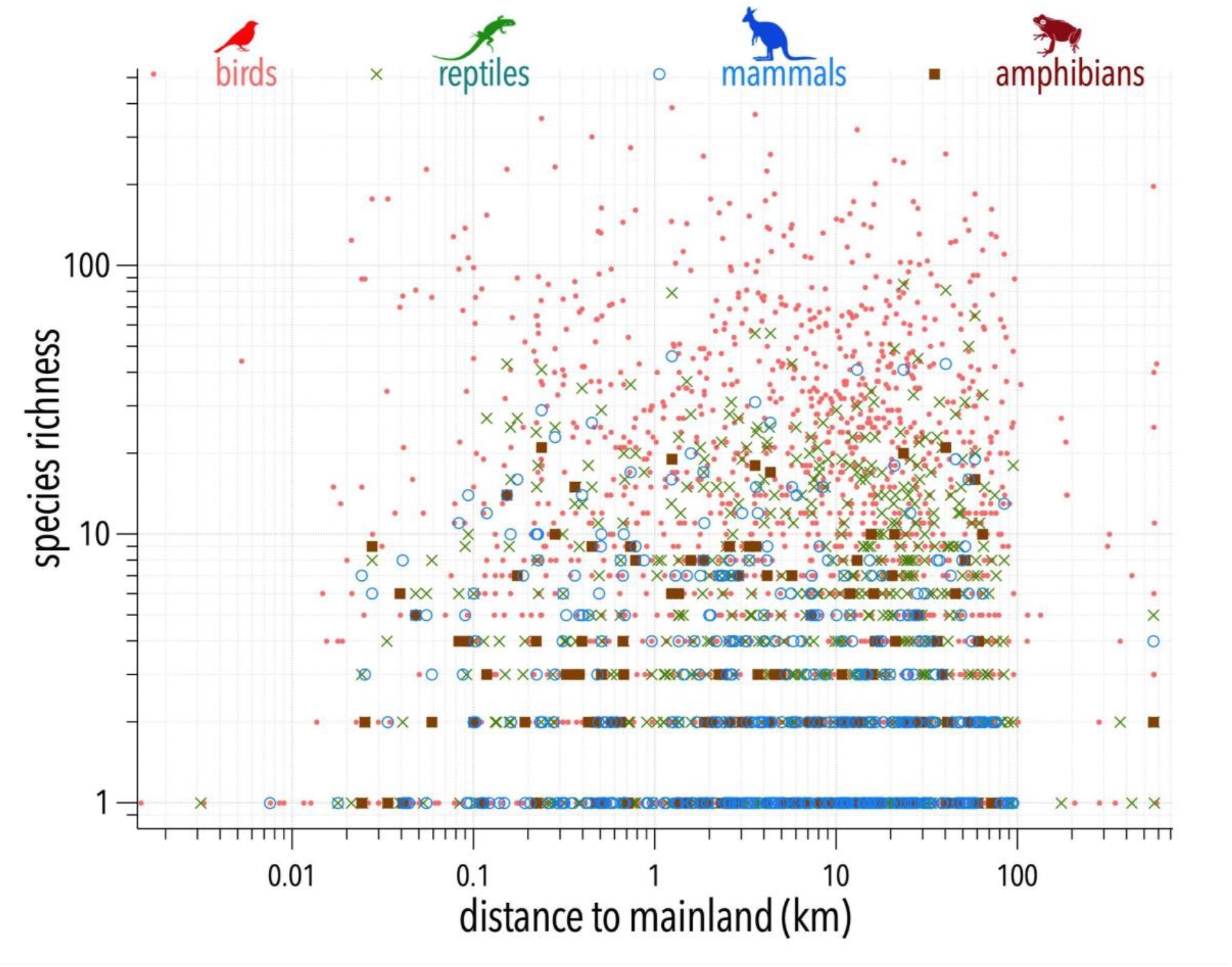
Power-law (log_10_-log_10_) relationship between species richness (log_10_ scale) and straight-line distance to the mainland (km; log_10_ scale) for birds, reptiles, mammals, and amphibians separately.

**Figure S8.**
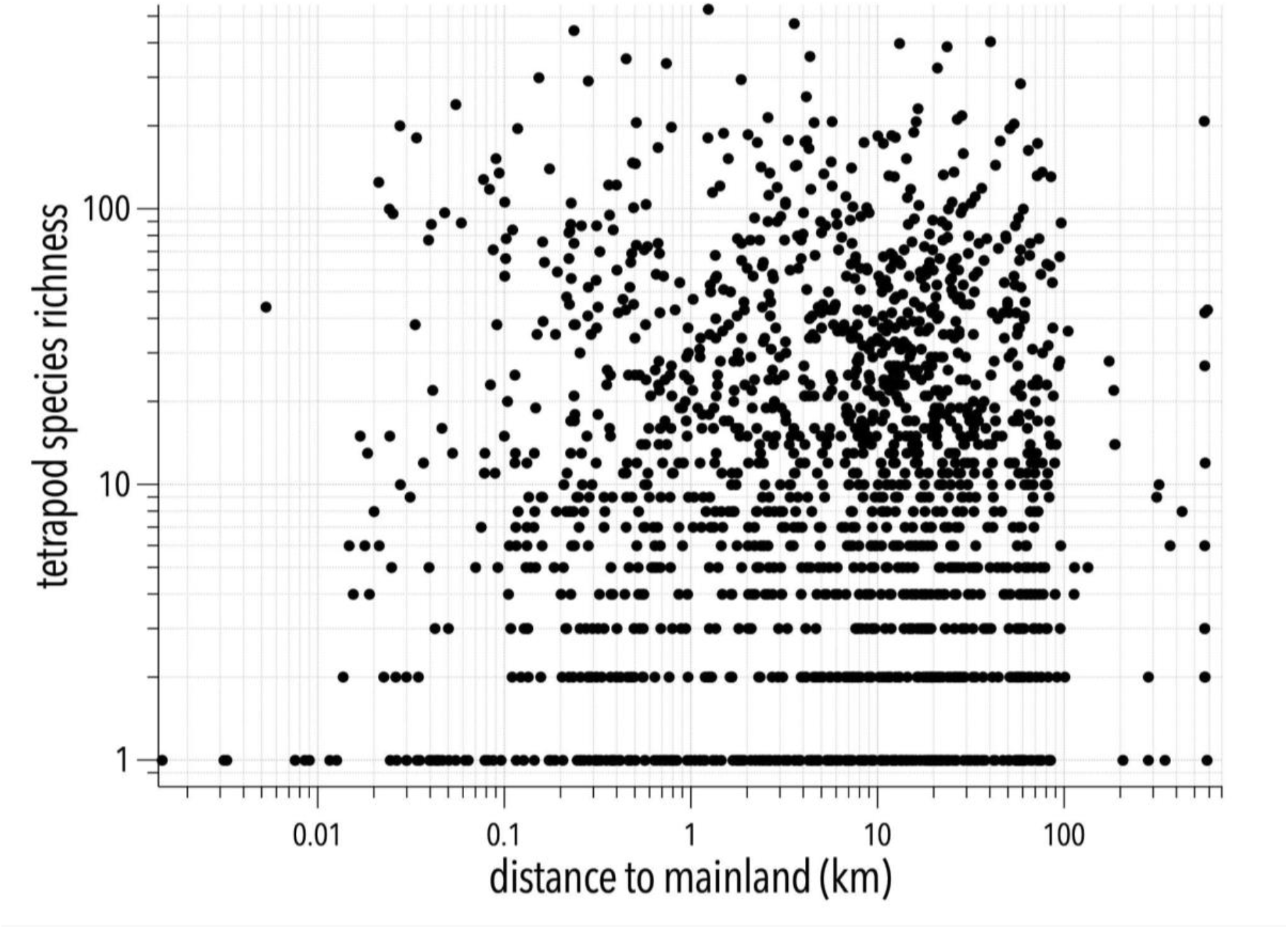
Power-law (log_10_-log_10_) relationship between species richness and straight-line distance to the mainland for terrestrial tetrapods combined..

**Figure S9.**
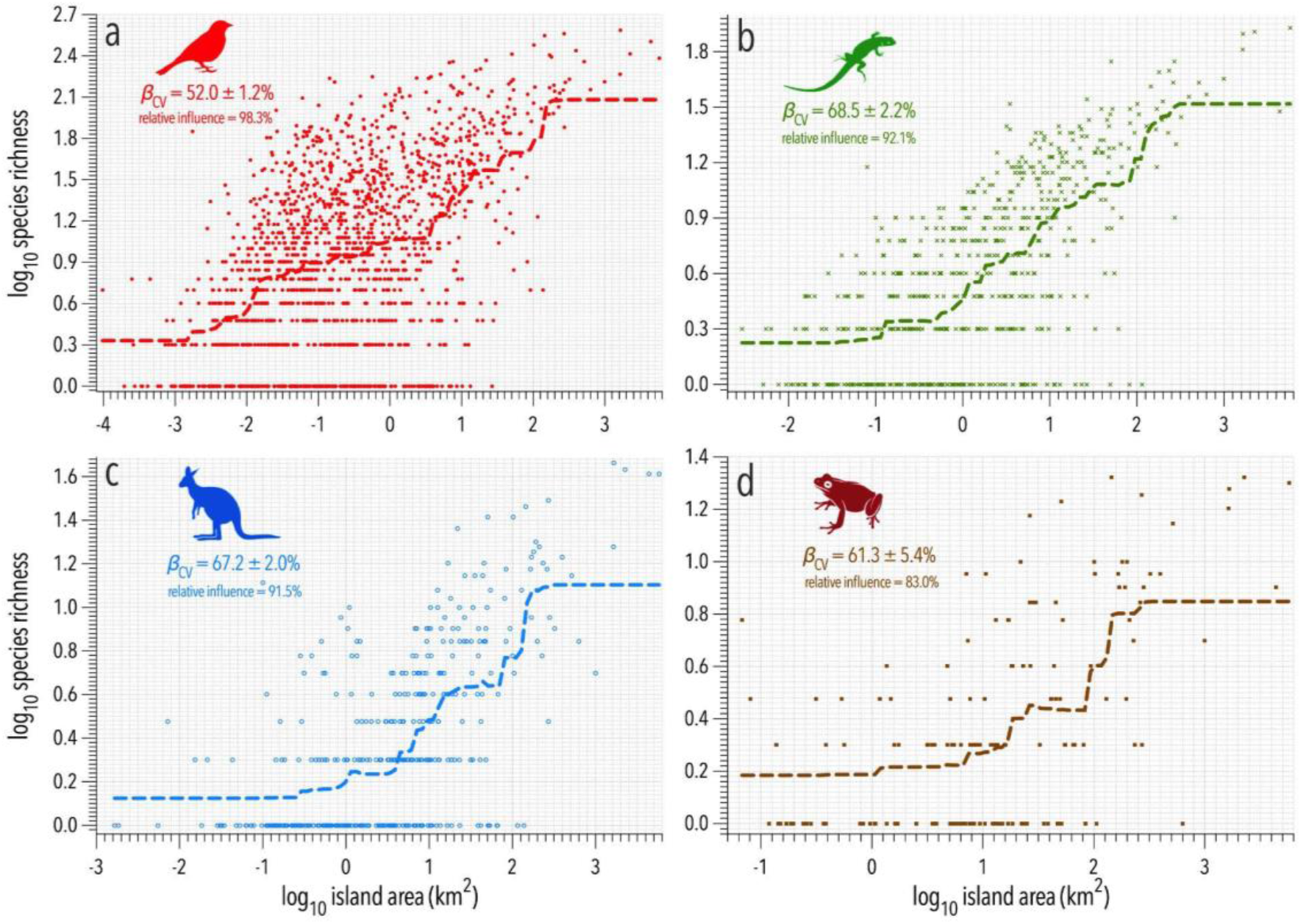
Boosted regression tree relationships between log_10_ species richness and log_10_ island area (km^2^) for each major tetrapod taxon separately. Shown in each panel is boosted regression tree coefficient of variation (*β*_CV_) as a measure of goodness of fit, as well as the relative influence of island area on the variance in species richness.

**Figure S10.**
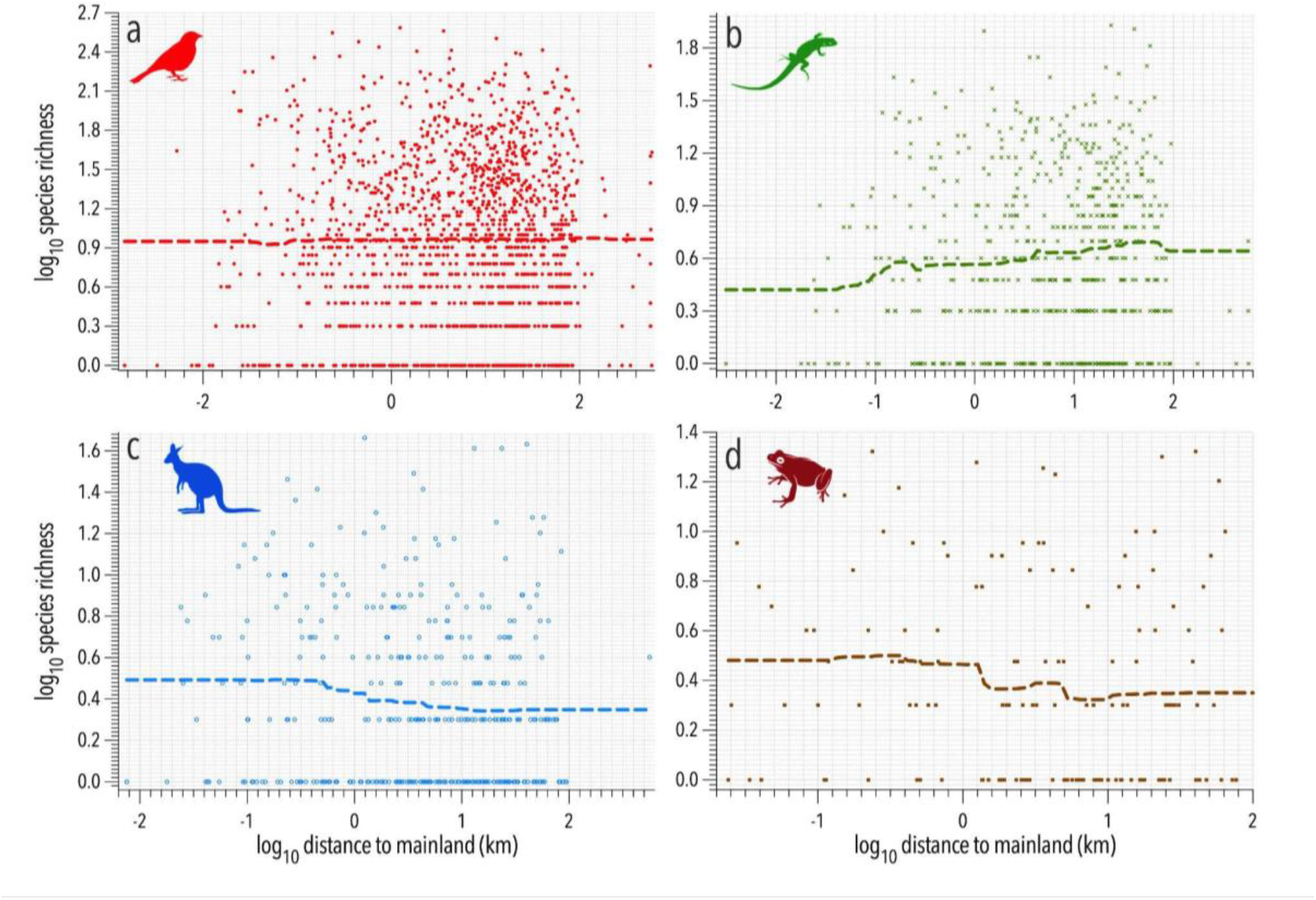
Boosted regression tree relationships between log_10_ species richness and log_10_ straight-line distance to mainland (km; isolation) for each major tetrapod taxon separately. Shown in each panel is boosted regression tree coefficient of variation (*β*_CV_) as a measure of goodness of fit, as well as the relative influence of island isolation on the variance in species richness.

**Figure S11.**
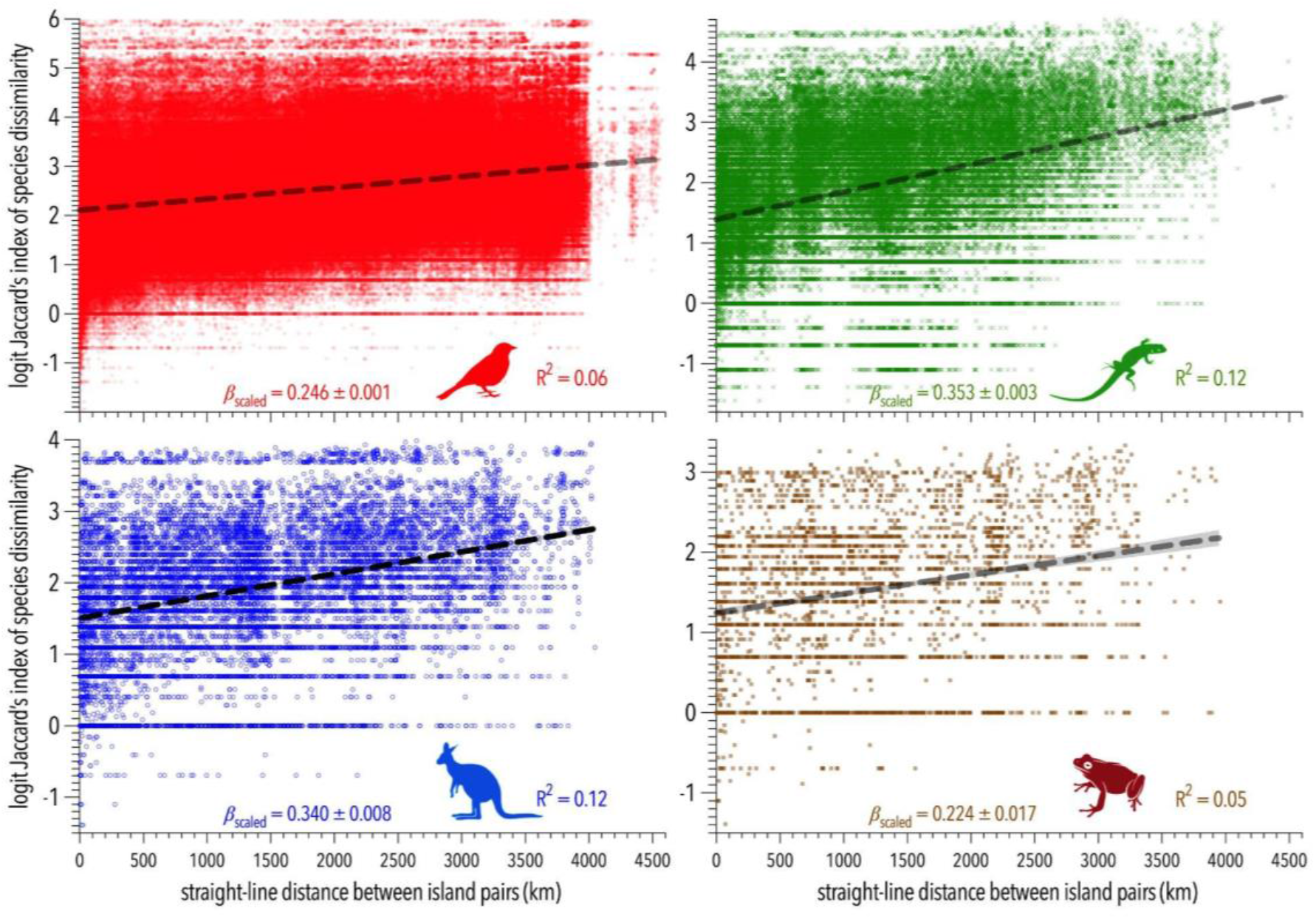
Jaccard’s index of species dissimilarity relative to straight-line distance (km) between islands for birds, reptiles, mammals, and amphibians.

**Figure S12.**
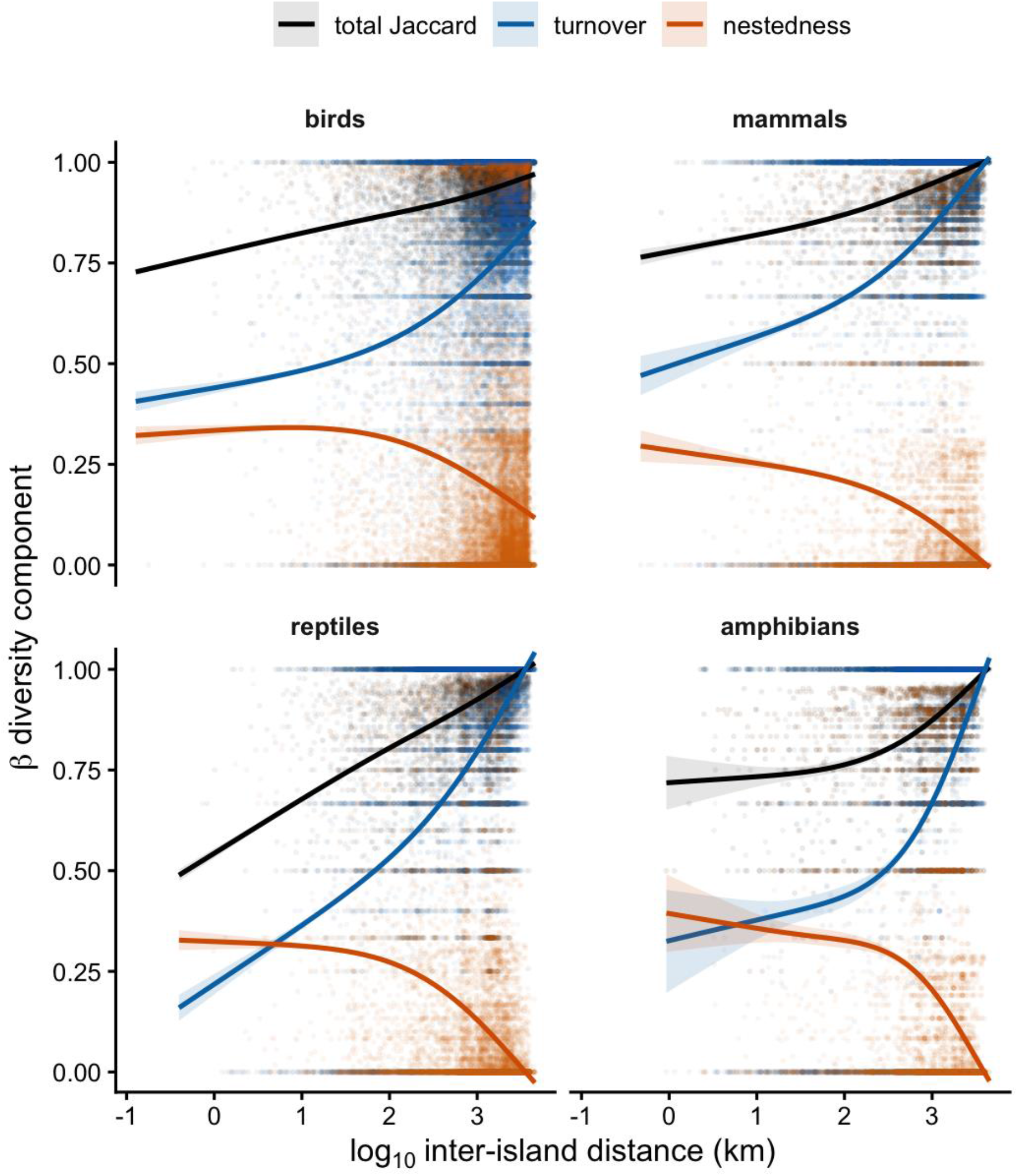
Total species dissimilarity and its turnover and nestedness-resultant components as functions of straight-line distance between island pairs for birds, mammals, reptiles, and amphibians. Components are shown together because turnover and nestedness are complementary parts of total Jaccard *β*-diversity. Increasing inter-island distance was associated primarily with increasing species turnover, whereas nestedness-resultant dissimilarity generally declined with distance, indicating that distant islands differed mainly through species replacement rather than ordered subset structure. Lines are separate linear summaries of each component and are not constrained to remain within the 0–1 bounds.

**Figure S13.**
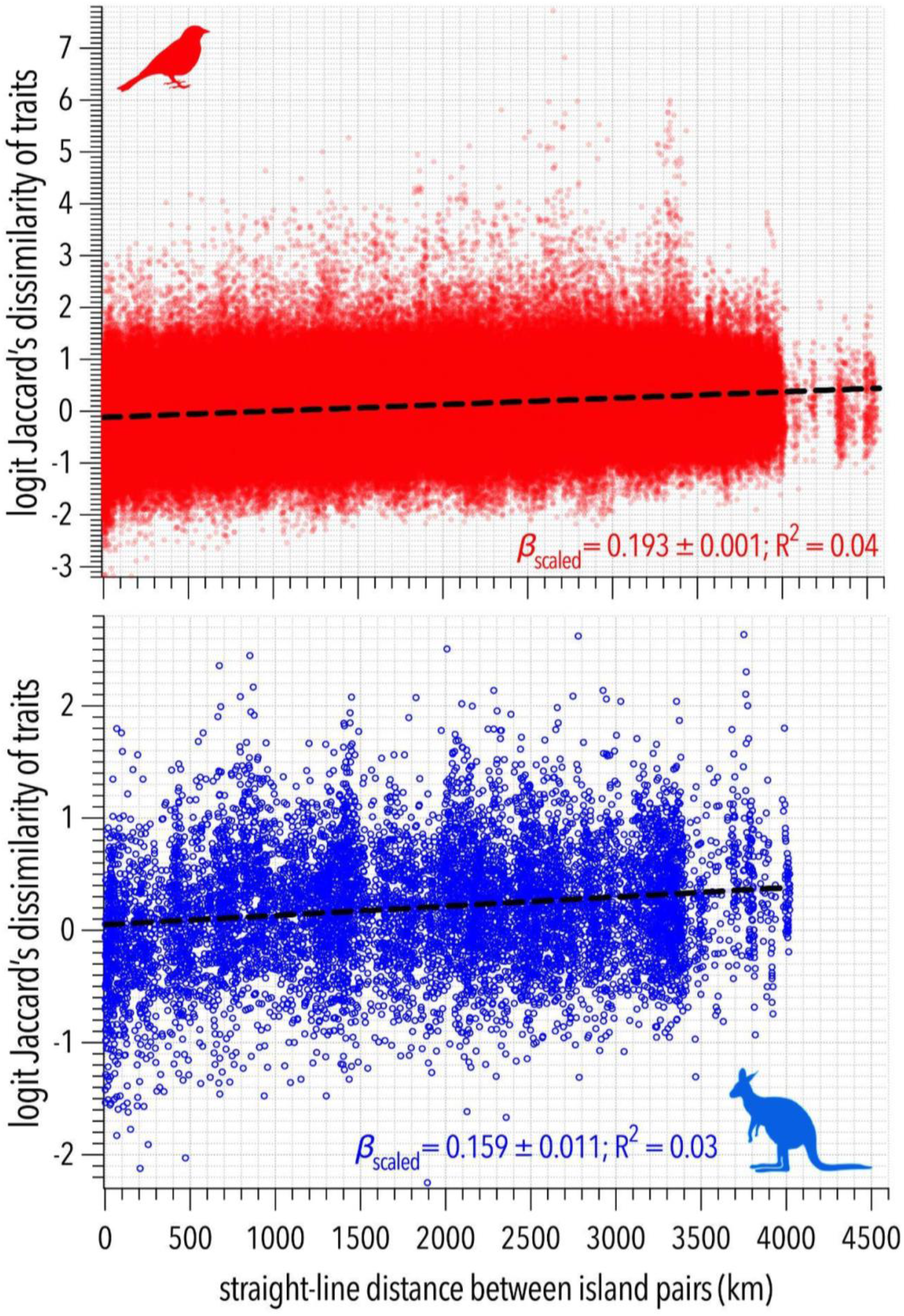
Jaccard’s index of trait dissimilarity (logit scale) relative to straight-line distance (km) between islands for birds (top panel) and mammals (bottom panel). Scaled linear fit parameters (slope = *β*_scaled_; goodness of fit = R^2^) shown for each taxon.

**Figure S14.**
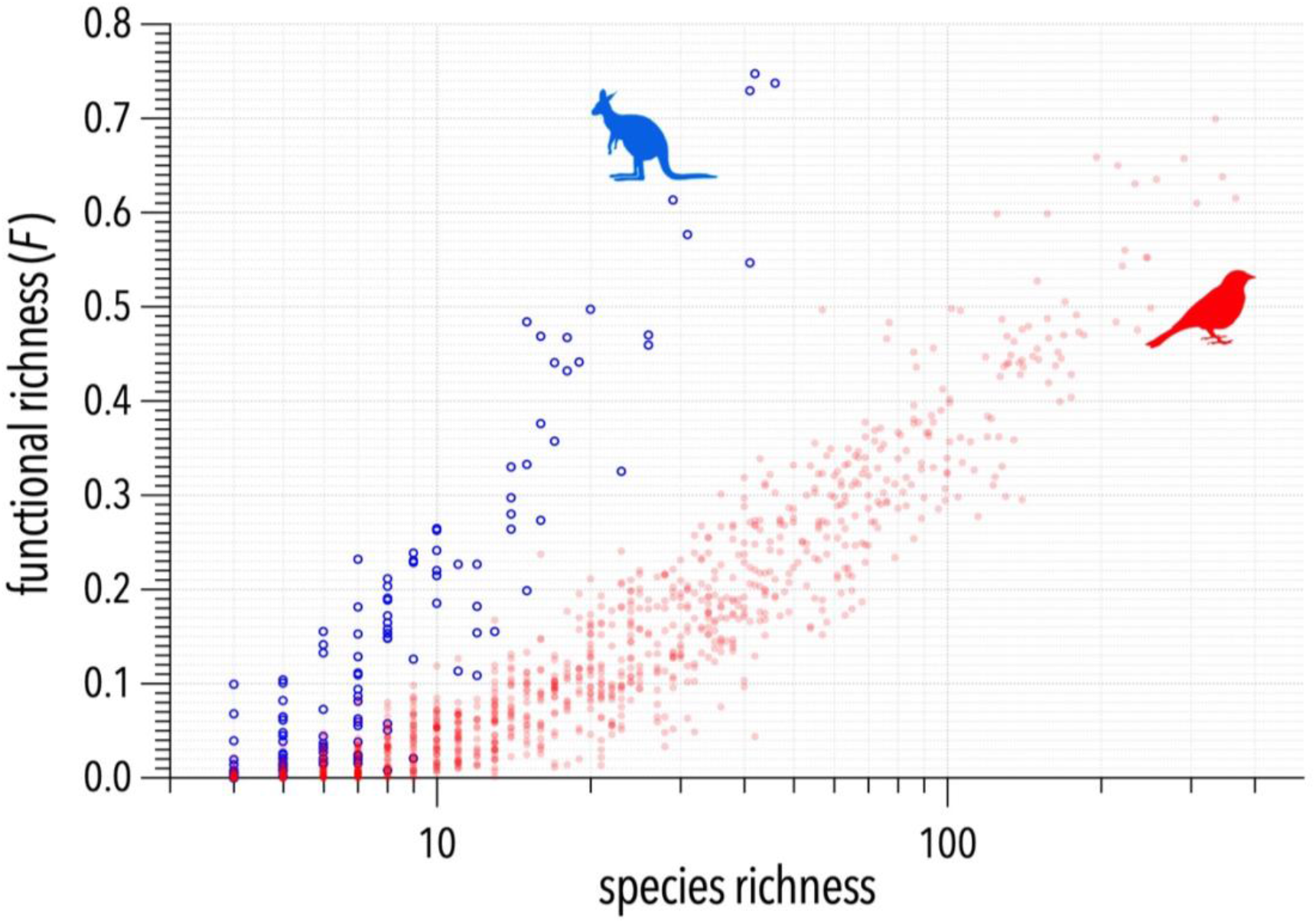
Relationship between functional richness and species richness (log_10_ scale) for mammals (blue) and birds (red).

**Figure S15.**
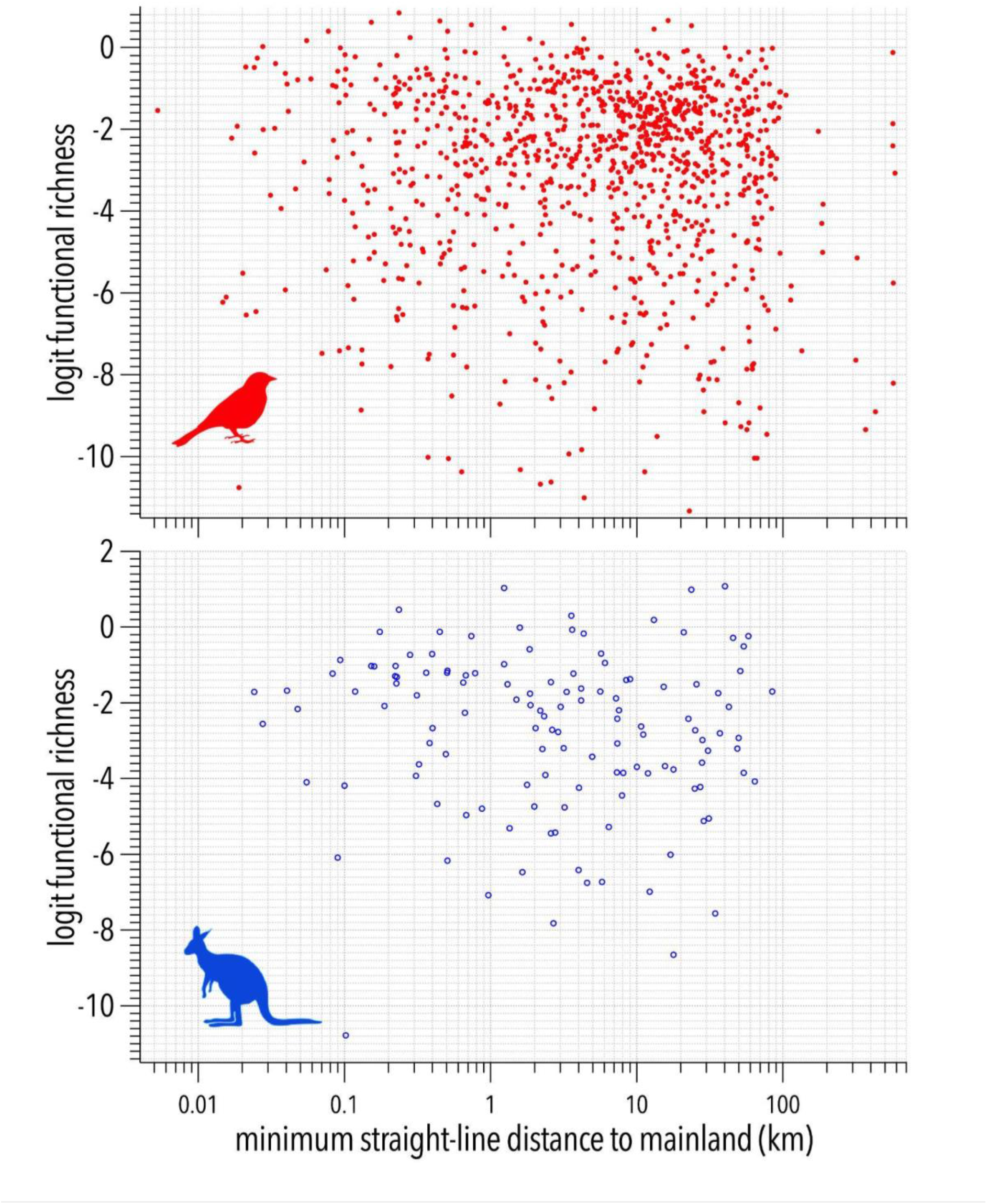
No relationship between logit functional richness and minimum straight-line distance between an island and the mainland (km; log_10_ scale) for bird (upper panel) and mammal (lower panel) traits.

**Figure S16.**
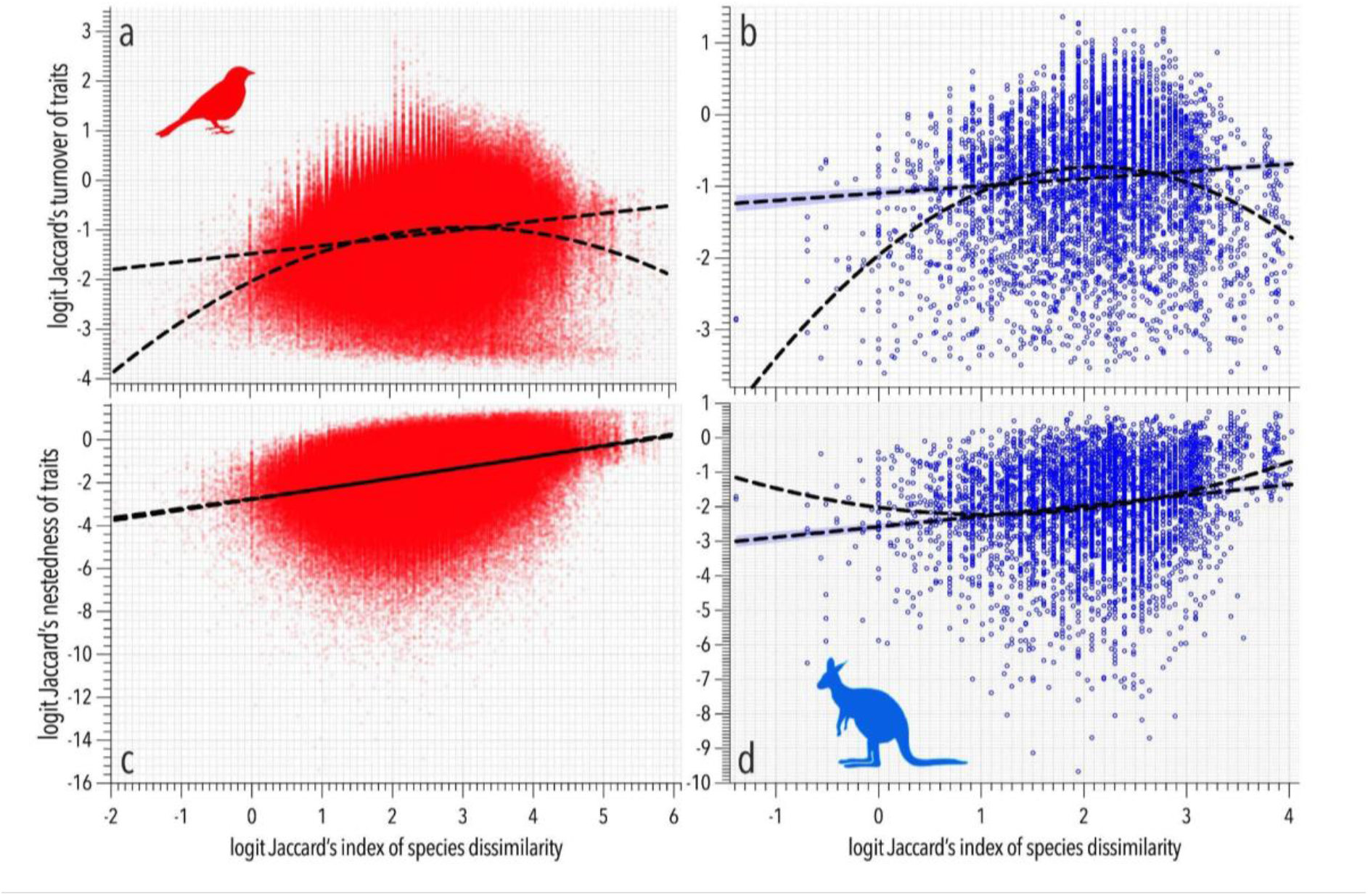
Trait turnover and nestedness relative to Jaccard’s index of species dissimilarity (logit scale). Relationship metrics using scaled *y* and *x* variables: (a) bird trait turnover: linear: R^2^ = 0.034, *β* = 0.206 ± 0.002; quadratic: R^2^ = 0.057; ER_q/l_ ≈∞; (b) mammal trait turnover: linear: R^2^ = 0.004, *β* = 0.105 ± 0.024; quadratic: R_2_ = 0.070; ER_q/l_ ≈∞; (c) bird trait nestedness: linear: R^2^ = 0.113, *β* = 0.316 ± 0.012; quadratic: R^2^ = 0.124; ER_q/l_ = 2.92; (d) mammal trait nestedness: linear: R^2^ = 0.028, *β* = 0.295 ± 0.025; quadratic: R^2^ = 0.038; ER_q/l_ = 1.38e^11^.

**Figure S17.**
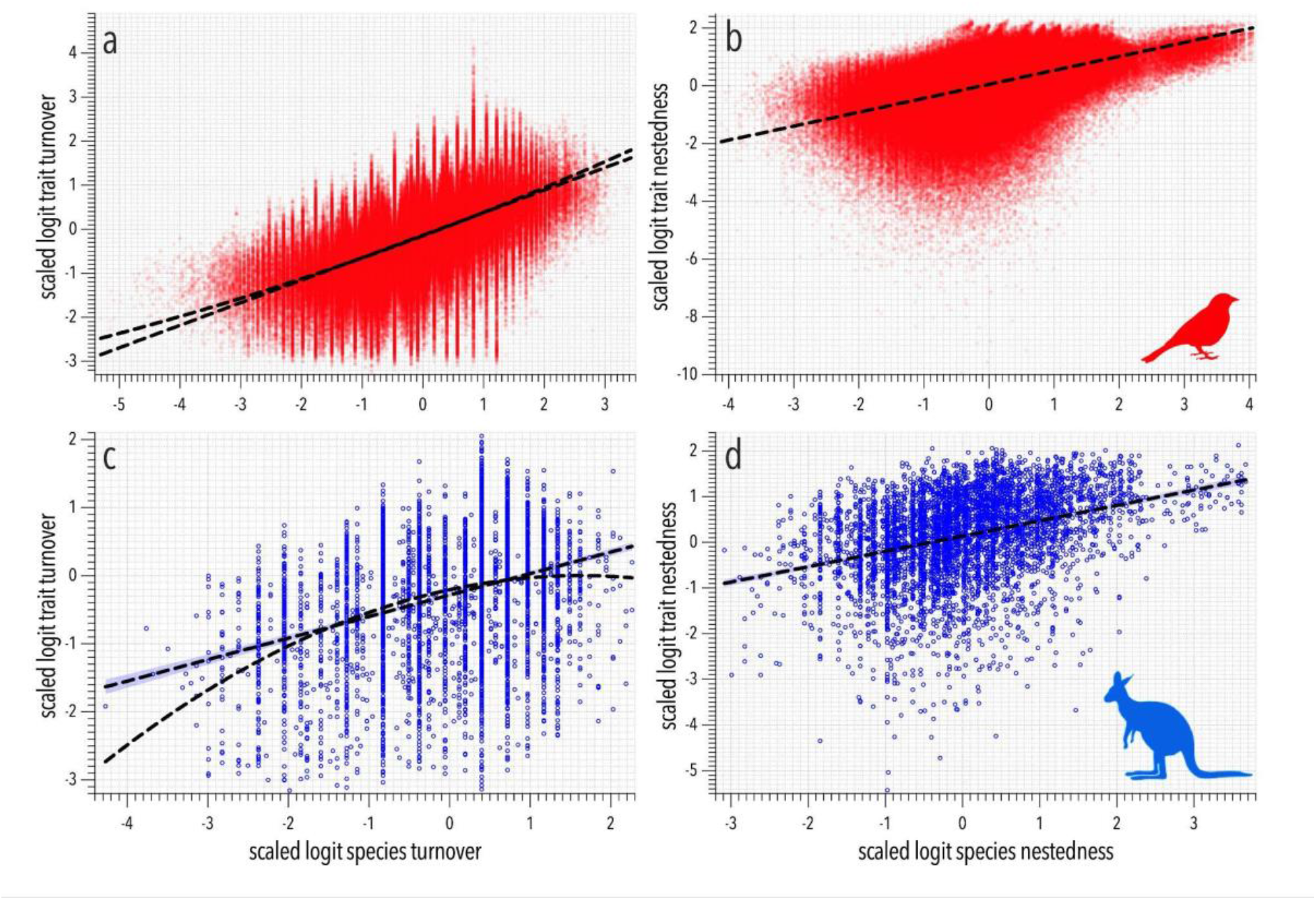
Trait turnover and nestedness relative to species turnover and nestedness (scaled logit). Relationship metrics using scaled *y* and *x* variables: (a) bird trait turnover *versus* species turnover: linear: R^2^ = 0.339, *β* = 0.511 ± 0.001; quadratic: R^2^ = 0.340; ER_q/l_ ≈ ∞; (b) bird trait nestedness *versus* species nestedness: R^2^ = 0.244, *β* = 0.483 ± 0.001; (c) mammal trait turnover *versus* species turnover: linear: R^2^ = 0.113, *β* = 0.316 ± 0.012; quadratic: R^2^ = 0.124; ER_q/l_ = 2.92; (d) mammal trait nestedness *versus* species nestedness: R^2^ = 0.116, *β* = 0.334 ± 0.013. There was more support (evidence ratio = 2.92) for a quadratic *versus* linear relationship in the expected direction (Fig. S17c). However, while there was also strong support for a quadratic relationship in birds (evidence ratio ≈ ∞), it was not in the expected direction, nor did it diverge much from the linear (Fig. S17a).

## Notes

https://doi.org/10.5281/zenodo.21736674

### Competing Interest Statement

The authors have declared no competing interest.

